# Exploring polymorphic interspecies structural variants in Eucalyptus: Unravelling Their Role in Reproductive Isolation and Adaptive Divergence

**DOI:** 10.1101/2023.10.20.563207

**Authors:** Scott Ferguson, Ashley Jones, Kevin Murray, Rose L. Andrew, Helen Bothwell, Benjamin Schwessinger, Justin Borevitz

## Abstract

Structural variants (SVs) play a significant role in speciation and adaptation in many species, yet few studies have explored the prevalence and impact of different categories of SVs. We conducted a comparative analysis of long-read assembled reference genomes of closely related *Eucalyptus* species to identify candidate SVs potentially influencing speciation and adaptation. Interspecies SVs can be either fixed differences, or polymorphic in one or both species. To describe SV patterns, we employed short-read whole-genome sequencing on over 600 individuals of *E. melliodora* and *E. sideroxylon*, along with recent high quality genome assemblies. We aligned reads and genotyped interspecies SVs predicted between species reference genomes. Our results revealed that 49,756 of 58,025 and 39,536 of 47,064 interspecies SVs could be typed with short reads, in *E. melliodora* and *E. sideroxylon* respectively. Focusing on inversions and translocations, symmetric SVs which are readily genotyped within both populations, 24 were found to be structural divergences, 2,623 structural polymorphisms, and 928 shared structural polymorphisms. We assessed the functional significance of fixed interspecies SVs by examining differences in estimated recombination rates and genetic differentiation between species, revealing a complex history of natural selection. Shared structural polymorphisms displayed enrichment of potentially adaptive genes. Understanding how different classes of genetic mutations contribute to genetic diversity and reproductive barriers is essential for understanding how organisms enhance fitness, adapt to changing environments, and diversify. Our findings reveal the prevalence of interspecies SVs and elucidate their role in genetic differentiation, adaptive evolution, and species divergence within and between populations.

## Introduction

Structural mutations that alter stretches of DNA greater than 50 bp in length have the potential to drastically change phenotypes (Alonge et al., 2020; Imprialou et al., 2017; Weischenfeldt et al., 2013) and contribute to population divergence and speciation (Marques et al., 2019; Zhang et al., 2021). Typically termed chromosomal rearrangements or structural variations (SV), these large mutations include inversions, translocations, duplications, insertions, and deletions (Savocco & Piazza, 2021). Until recently however, technological constraints, namely sequencing read lengths, have inhibited their discovery (Sedlazeck et al., 2018), and their role in population evolutionary processes remains poorly understood (Pokrovac & Pezer, 2022). Using third-generation long-read sequencing, such as those offered by Oxford Nanopore Technologies and PacBio, evolutionary genomic studies can now affordably assemble highly contiguous genomes of several individuals across related species. The next challenge is to perform population scale SV discovery and examine the role of SVs in population divergence and speciation.

Structural variation can occur in all parts of the genome: coding, noncoding, and repetitive regions such as transposons, telomeres and centromeres. When they occur within coding regions, they may alter regulatory elements, introns, exons, whole genes, or multiple genes (Radke & Lee, 2015; Stewart & Rogers, 2019). Even when they do not occur within coding regions they can change the chromatin structure and impact gene expression (Kim et al., 2019; Shanta et al., 2020). Different SV types are known or predicted to have different genomic effects. Inversions can inhibit recombination between different arrangements, reducing the overall recombination rates between homologous chromosome pairs, and fixing the alleles captured within their bounds (Thompson & Jiggins, 2014). Inversion-linked, cosegregating alleles can become reproductively isolated and purged through underdominant selection, due to increased sterility of heterozygous individuals (Kirkpatrick & Barton, 2006; Lande, 1985; Walsh, 1982). However, a novel inversion, if adaptive, may provide enough selective advantages to outweigh its disadvantages, be selected for, and rise to high frequency within populations (Harringmeyer & Hoekstra, 2022; Rieseberg, 2001). Translocations, while less studied than other rearrangements (Robberecht et al., 2013), may have similar genomic effects as inversions (Ortiz-Barrientos et al., 2016). Duplications, highly common and also likely to be selected against (Flagel & Wendel, 2009; Wu & Cox, 2019), could be preserved due to their ability to acquire new function (neofunctionalisation) or by retaining a subset of original function (subfunctionalisation) (Braasch et al., 2016; Freeling et al., 2015; Lien et al., 2016; Wu & Cox, 2019). Large (> 50 bp) insertions and deletions, which are often genotyped as presence/absence variants (PAVs), copy number variations (CNV), or gene duplications, are also very common within genomes (Conrad & Hurles, 2007; Pokrovac & Pezer, 2022). These SVs are known to impact genes and gene structure, and to affect phenotypes (Sun et al., 2022; Yuan et al., 2021), although many can also be neutral.

An ancestral population, once highly syntenic, undergoes division into two non-interbreeding groups, with structural variations (SVs) emerging between them, Figure 1. These interspecies SVs can be genotyped as fixed within one species, leading to structural divergence (SD) or polymorphic within one species, termed structural polymorphisms (SP) (Ferguson, et al., 2022). Adding complexity, SVs can also be genotyped as polymorphic in both populations, referred to as shared structural polymorphisms (SSP). To classify interspecies SVs, genotyping within both species is essential, enabling us to categorise them based on their presence/absence in population 1 and population 2 as fixed/absent (SD), fixed/polymorphic (SP), absent/polymorphic (SP), or polymorphic/polymorphic (SSP). The rate at which SVs are SD, SP, or SSP is unknown; however, rates will depend on the evolutionary distance between populations or species, effective population size, and mutation rate, among other factors. If the status of an SV remains uncertain, inferences of its impact on divergence and adaptation are difficult.

**Figure 1.**
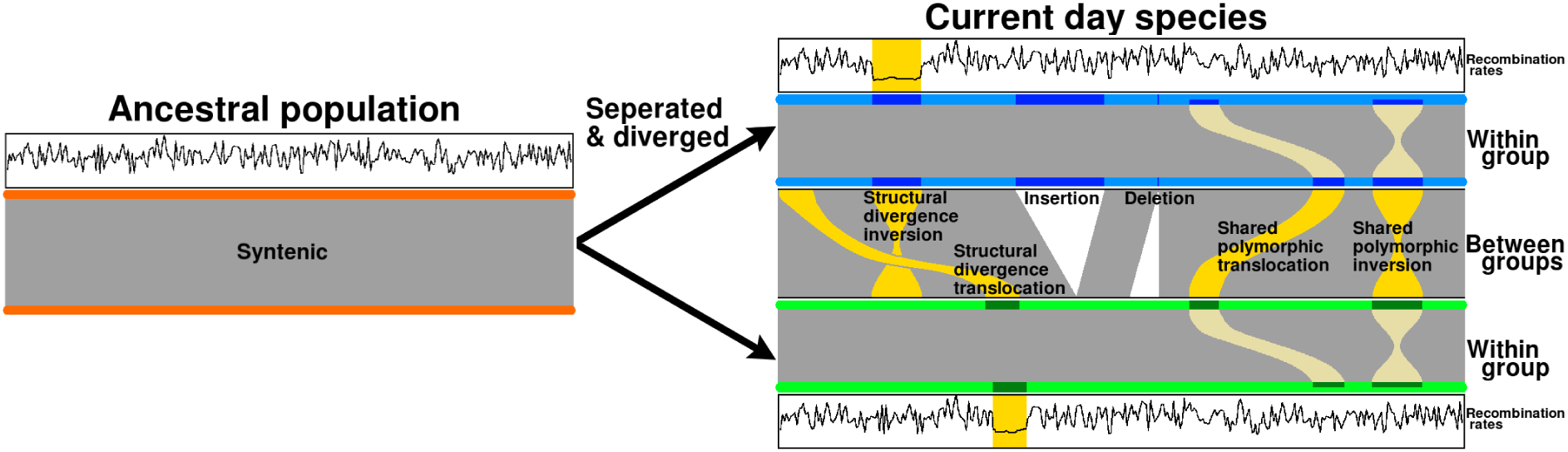
Structural variations within sister species. The once highly syntenic ancestral population separates and divides into two non-interbreeding groups. Structural variations, which reduce genome-wide synteny, discovered between the two groups may be genotyped within populations as fixed or polymorphic. When fixed in a single population, SVs become a structural divergence (SD). If polymorphic within one population, SVs become structural polymorphisms (SP), or if polymorphic in both populations a shared structural polymorphisms (SSP). The different classes of population genotyped SVs may have different impacts on recombination rates, divergence, and adaptation.

Analysing the genomic differences between recently diverged species has revealed genome regions involved in reproductive isolation (Hejase et al., 2020), adaptive genes (Eshel et al., 2021), and the genome-wide landscape of diversification between and within chromosomes (Henderson & Brelsford, 2020; Piatkowski et al., 2023; Zhang et al., 2022). Here using two closely related *Eucalyptus* species, *E. melliodora* and *E. sideroxylon* (Ferguson et al., 2023; Thornhill et al., 2019), we genotype SVs within their respective populations and calculate their rates of population variability. Structural variation rates are compared to find evidence of SD, SP, and SSP. Additionally, we examine recombination rates and Fixation Index (*F_ST_*) within population fixed SVs to assess allele fixation and accelerated evolution between populations.

## Methods

### Population sampling and sequencing

Yellow box (*Eucalyptus melliodora*) and red ironbark (*E. sideroxylon*) are closely-related eucalypts of the box-gum grassy woodland endangered ecological community. These species are often found growing in sympatry or parapatry, and widely hybridise throughout their ranges in southeastern Australia. We collected 472 *E. melliodora* and 180 *E. sideroxylon*, all samples being wild and undomesticated. Samples were environmentally stratified to capture major clines in climate-adaptive genomic variation across the species’ distributions. GPS data was recorded for each sample, and leaf material was dried in silica desiccant (Figure 4).

Twenty 3 mm disc punches (UniCore, Qiagen) from each leaf sample were placed in mini-tubes with a 3 mm ball bearing, frozen with liquid nitrogen and ground in a TissueLyser II (Qiagen). Genomic DNA was extracted using a 96-well plate column-based kit; Stratec Invisorb DNA Plant HTS 96 Kit/ C, according to the manufacturer’s instructions (Stratec SE, Birkenfeld, Germany). DNA was quantified using a Infinite M1000 PRO Tecan fluorescence microplate reader (Tecan Trading AG, Switzerland), and standardised to 1 ng/µL, using a liquid handling robot. Library preparation was performed using a modified Illumina Nextera DNA Library Prep Kit workflow, which is available in Protocols.io and described in Jones et al (2023). Libraries were then quantified using GXII and Quant-iT, and pooled for equal representation. Prior to size selection, samples were concentrated using 2x binding buffer and 100 µL of Sera-Mag Speedbeads Carboxylate-Modified Particles (Thermo Scientific, Fremont, CA, USA). Size selection was then performed on a Pippin Prep (Sage Science, Inc., Beverly, MA, USA), for 400-650 bp fragments. Samples were again concentrated with 2x binding buffer and 100 µL of Sera-Mag beads, then quantified using both a Qubit Fluorometer (Thermo Scientific, Fremont, CA, USA) and Bioanalyzer high sensitivity DNA chips (Agilent Technologies, Santa Clara, CA, USA). Whole genome sequencing was performed on an Illumina NovaSeq 6000, 150 bp paired end sequencing, by Novogene (HK) Co., Ltd (Hong Kong).

### Genome scaffolding

We performed Hi-C scaffolding, grouping, ordering, and orienting of our previously assembled *E. melliodora* genome into pseudo-chromosomes (Ferguson, et al., 2022). Leaves were obtained from the reference tree, and a proximity ligation library for chromosome conformation capture was created with a Phase Genomics Proximo Hi-C (Plant) Kit (version 4), according to the manufacturer’s instructions (document KT3040B). DpnII, HinFI, MseI, DdeI to digest the genome. Sequencing was performed on an Illumina NovaSeq 6000, 150 bp paired end sequencing. Hi-C scaffolding began by aligning all Hi-C reads to *E. melliodora’s* contigs using bwa mem (version: 0.7.17; parameters: −5SP; Li, 2013). Next, PCR duplicates were identified with Samblaster (Faust & Hall, 2014) (version: 0.1.26). Linkage information captured within Hi-C reads was assessed with Juicer (version: 1.6; Durand, Shamim, et al., 2016) and scaffolding was performed using 3D-DNA (version: 190716; parameter: −i 1000; Dudchenko et al., 2017). Due to the high repeat content, Hi-C read coverage was highly variable and resulted in poor quality scaffolding. To account for variability in read coverage, we ran 3D-DNA with “--editor-repeat-coverage 5”, altering the misjoin detection threshold. After initial scaffolding the Hi-C contact map was manually edited with Juicebox (version: 2.16; Durand et al., 2016). Previously assembled contigs for *E. sideroxylon* (Ferguson et al., 2022) were scaffolded with RagTag (version: v2.1.0; Alonge et al., 2022) using synteny to our Hi-C scaffolded *E. melliodora* genome.

Genome completeness was measured with BUSCO (version 5; Manni et al., 2021) and long terminal repeat assembly index (LAI; Ou et al., 2018). BUSCO scores genome completeness by identifying and reporting on the proportion of lineage specific highly conserved single-copy genes; more complete genomes have a high proportion of identified BUSCO genes. LAI identifies long terminal repeat (LTR) sequences and reports on the proportion that are intact; more complete genomes have a high proportion of intact LTR sequences.

### Genome annotation

Genomes were annotated for transposable elements (TE) using genome-specific, *de novo* repeat libraries created with EDTA (version: 1.9.6; Ou et al., 2019) and RepeatMasker (version: 4.1.1; Smit et al., 2020). RepeatMasker additionally annotated our genomes for simple repeats. Repeat masked genomes were next annotated for genes using BRAKER2 (version 2.1.6; Brůna et al., 2021). BRAKER2 was run with ProtHint (version2.6.0; Brůna et al., 2020) and GeneMark-EP (version: 4; Brůna et al., 2020). ProtHint analysed training proteins to determine their evolutionary distance to the genome, aiding GeneMark-EP to train a gene detection model. Training protein sequences were obtained from the National Center for Biotechnology Information (NCBI; Sayers et al., 2021) and included all available transcripts for Myrtaceae (Taxonomy ID: 3931) and *Arabidopsis thaliana* (Taxonomy ID: 3702).

Candidate genes were functionally annotated for eggNOG orthogroup, COG category, GO term, KEGG term, and PFAM using eggNOG-mapper (version: 2.1.12; parameters: −m diamond --itype CDS --tax_scope Viridiplantae; Cantalapiedra et al., 2021).

### Synteny and structural variation annotation

Shared sequences were identified between genomes by alignment with nucmer (parameters: --maxmatch -l 40 -b 500 -c 200), from the MUMmer (version: 3.23; Kurtz et al., 2004) toolset. Nucmer identifies all shared 40-mers between the two genomes and joins all 40-mers within 500 bp into single alignments. After aligning the two genomes MUMmer’s delta-filter (parameters: -i 80 -l 200) tool removes all alignments < 200 bp and with an identity < 80%. A low sequence identity score (80%) was used due to the high heterozygosity of *Eucalyptus* genomes (Murray et al., 2019), and a higher score may incorrectly filter out real alignments. Using SyRI (version: 1.5; Goel et al., 2019), filtered nucmer alignments were analysed and subsequently genomes were annotated for synteny, inverted, translocated, duplicated, and not-alignable regions.

All inversions, translocations, duplications, and unaligned regions described by SyRI were genotyped for all 563 samples within both species using Paragraph (Chen et al., 2019) and our short-read alignments.

A 0/1/2 matrix was created for all genotyped SV within both species and for all categories of SV. Using the R (R Core Team, 2023) function Cor, the correlation between SVs of interest was calculated and visualised with heatmap.

### Alignment and variant calling

Raw population sequences were trimmed (sequencing adaptors and barcodes), quality filtered (average quality score < 20), and merged (overlapping read pairs were combined into single reads) using AdapterRemoval (version: 2.3.0; Schubert et al., 2016). Genome coverage was estimated for each sample and samples with low coverage (< 10x) were removed. Quality filtered reads were next aligned to both reference genomes (*E. melliodora* and *E. sideroxylon*) using bwa mem (parameters: -p). Samples with <75% alignment were then removed. Aligned reads for all remaining samples were variant called with bcftools (version: 1.12; Danecek et al., 2021) mpileup (parameters: MAPQ > 30, base quality > 15). The default mutation rate (0.0011) was increased to 0.01, making variant calling more robust when calling low coverage heterozygous SNPs. Variant files were then merged, resulting in four datasets; (reference genome - population species) *E. melliodora - E. melliodora*, *E. melliodora - E. sideroxylon*, *E. sideroxylon - E. melliodora*, and *E. sideroxylon - E. sideroxylon*.

### Variant filtering

Using bcftools norm, multiallelic variants for each variant dataset were decomposed into multiple single variants. Decomposed variants were filtered, removing variants present in < 10% of samples and with less than 20 supporting reads, within each dataset using bcftools view. Variants were next recomposed, all remaining multiallelic variants rejoined, and each dataset further filtered to remove all indels and multiallelic SNPs (Murray, 2023).

High-quality, biallelic SNP datasets for each reference genome were combined, and a principal component analysis (PCA) performed with PCAngsd (version: 1.10; Meisner & Albrechtsen, 2018). Visual inspection of PCA plots allowed identification and removal of hybrids, outliers, and incorrectly labelled samples.

### SNP phasing and recombination calculation

Before computing ρ (estimated recombination rate) within our four datasets, SNPs first required phasing. Phasing links each variant allele, placing them into haplotype blocks, separating maternal and paternal variants. As the linkage information provided by paired-end short reads is not capable of phasing all SNPs, a two-step phasing process was used. First, individual samples were extracted from species variant files into a single sample variant file and using read alignments, SNPs, when possible, were phased with WhatsHap (version: 1.7; Martin et al., 2016). Second, partially phased sample variant files were re-merged and the Hidden Markov Model (HMM) phaser SHAPEIT4 (version: 4.2.2; Delaneau et al., 2019) inferred haplotypes and phased the remaining unphased SNPs. Parameters (--use-PS 0.0001 --mcmc-iterations 6b,1p,1b,1p,1b,1p,1b,1p,8m --pbwt-depth 6 --sequencing) specified for SHAPEIT4 were optimised by balancing maximum accuracy and runtime. At the completion of this two-stage phasing approach all SNPs for each dataset were phased. After phasing, ρ was calculated for each dataset using LDJump (parameters: alpha = 0.05; version: 0.3.1; Hermann et al., 2019), specifying a window size of 1 kb. LDJump made use of LDHat (version: 2.2a; Auton & McVean, 2007) to decrease runtime.

### Fixation index (***F_ST_***)

Filtered SNP datasets were combined for each reference genome, and subsequently *F_ST_* was calculated for each SNP using PLINK (version: 1.9; Chang et al., 2015). Per SNP *F_ST_* values were averaged for each region of interest for further analysis.

## Results

### Genome scaffolding

We generated Hi-C data and performed Hi-C scaffolding to order, orient, and combine contigs into pseudo-chromosomes for *E. melliodora*. Hi-C sequencing generated 45.48 Gbp in 151,590,503 paired reads, giving an estimated genome coverage of 71.14x. After aligning Hi-C reads to *E. melliodora’s* contigs and identifying PCR duplicates, 18,507,548 (12.21%) read pairs were found to contain linkage information. Further examination showed that 9,612,532 (6.34%) read pairs spanned contigs, and 8,895,016 (5.87%) read pairs were contained within a single contig. Non-informative reads were either chimeric, unmapped, PCR duplicates, or had low mapping quality (MAPQ < 30, mostly due to multi-mapping of short reads to repeat regions). For all Hi-C statistics see Supplementary Table S1. Using 3D-DNA *E. melliodora’s* contigs were scaffolded (Supplementary Figures S1 and S2). Contigs for *E. sideroxylon* were syntenically scaffolded against *E. melliodora’s* Hi-C scaffolded genome. Both BUSCO and LAI scores indicate that both genomes are highly complete (Table 1).

**Table 1.** Genome assembly statistics for *E. melliodora* and *E. sideroxylon*.

|  | *E. melliodora* | *E. sideroxylon* |
| --- | --- | --- |
| **Scaffolded genome Size (bp)** | 639,266,298 | 592,154,182 |
| **% of genome in scaffolds** | 97.60% | 98.15% |
| **Scaffold N50 (Mbp)** | 59.47 | 60.48 |
| **Contig N50 (Mbp)** | 1.87 | 5.22 |
| **Contig count** | 564 | 297 |
| **BUSCO complete** | 98.54% | 96.47% |
| **LAI** | 18.31 | 18.70 |
| **Repetitive % (TE %)** | 48.50% (47.13%) | 47.83% (46.58%) |
| **Gene candidates** | 58,902 | 57,299 |
| **Proportion of genome in gene candidates** | 21.85% | 21.04% |

### Annotation (repeats & genes)

Both genomes were annotated for transposable elements (TE), simple repeats, and genes (Table 1). Transposable elements and simple repeats were annotated with genome-specific *de novo* repeat libraries. Soft repeat masked genomes were next annotated for genes.

### Synteny and structural variation annotation

Shared sequences between *E. melliodora* and *E. sideroxylon* were identified using nucmer from the MUMmer toolset. Subsequently, using SyRI, shared sequences were classified as syntenic, inverted, translocated, or duplicated, and both genomes accordingly annotated for these regions. Additionally, both genomes were annotated for unaligned regions, which are unique to each genome, resulting from insertions, deletions, or divergence beyond recognition. 85.94% of *E. melliodora’s* genome was found to be shared with *E. sideroxylon’s* genome; conversely, 87.70% of *E. sideroxylon’s* genome was found to be shared with *E. melliodora’s* genome. The majority of shared sequences were syntenic. A more detailed analysis of alignment types showed that syntenic regions are, on average, frequent and large, inversions are rare and typically very large, translocations are moderately sized and frequent, duplications are very frequent and small, and unaligned regions are very frequent and small (Table 2, Figure 2).

**Figure 2.**
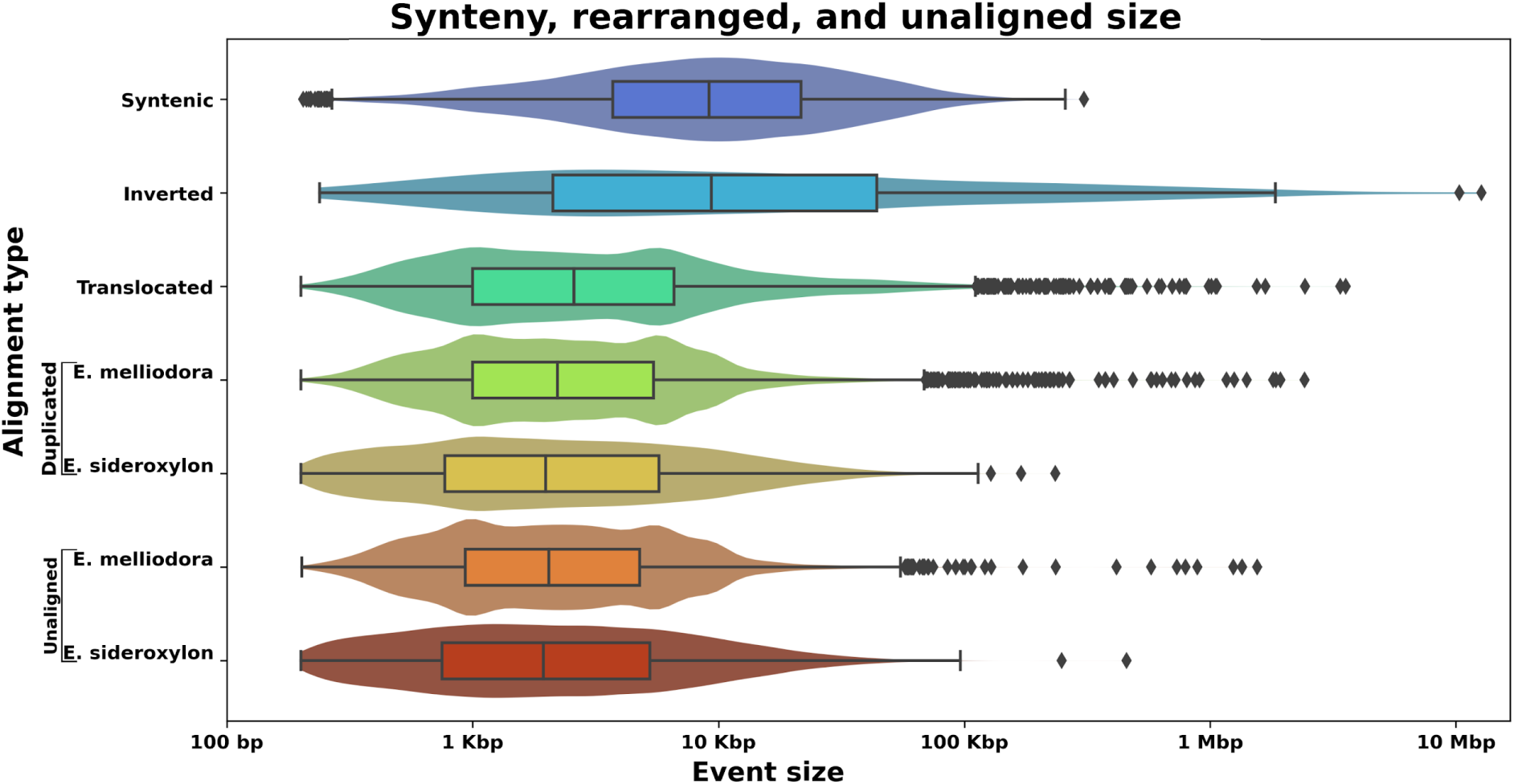
Synteny, rearranged, and unaligned event sizes. As syntenic, inverted, and translocated regions are approximately the same size with each genome (differing only by small indels) these alignment types are only shown for *E. melliodora*. Duplications and unaligned regions are unique to each genome and as such are shown for both *E. melliodora* and *E. sideroxylon*. See Supplementary Figure S3 for all event sizes for both genomes.

**Table 2.** Proportion, number of regions, and total amount the genome that was found to be syntenic, rearranged, and unaligned within *E. melliodora* and *E. sideroxylon* when their genomes were aligned.

| Genome | Statistic | Syntenic | Inversion | Translocation | Duplication | Unaligned |
| --- | --- | --- | --- | --- | --- | --- |
| *E. melliodora* | Count | 19,137 | 232 | 10,645 | 26,762 | 20,386 |
|  | Average size (Kbp) | 16.18<br>± 20.93 | 202.96<br>± 1097.87 | 11.41<br>± 69.65 | 5.25<br>± 32.57 | 4.30<br>± 7.13 |
|  | Total (Mbp) | 309.60 | 47.09 | 121.49 | 140.63 | 87.74 |
|  | Proportion | 49.62% | 7.55% | 19.47% | 22.54% | 14.06% |
| *E. sideroxylon* | Count | 19,137 | 232 | 10,645 | 20,102 | 18,777 |
|  | Average size (Kbp) | 16.14<br>± 20.87 | 177.67<br>± 851.99 | 11.29<br>± 65.34 | 4.30<br>± 33.33 | 3.81<br>± 6.67 |
|  | Total (Mbp) | 308.78 | 41.22 | 120.22 | 86.44 | 71.51 |
|  | Proportion | 53.13% | 7.09% | 20.69% | 14.87% | 12.30% |

### Variant calling and PCA

After calculating the total number of sequenced bases per sample and removing samples which had low coverage (< 10x), *E. melliodora’s* samples yielded on average 9.49 Gbp (range: 6.27 Gbp - 27.22 Gbp). Similarly, *E. sideroxylon’s* samples yielded on average 9.10 Gbp (range: 5.82 Gbp - 28.87 Gbp). Examined across both populations and both reference genomes, coverage averaged 15.40x (range: 10.00x - 48.7x). After aligning both populations sequences to both reference genomes, and filtering out samples with low alignment (< 75%), an average of 96.55% (range: 77.91% - 98.80%) of reads aligned to both genomes. Variants were called for the remaining samples, resulting in four dataset; (reference genome - population species) *E. melliodora - E. melliodora*, *E. melliodora - E. sideroxylon*, *E. sideroxylon - E. melliodora*, and *E. sideroxylon - E. sideroxylon* (Table 3, Figure 3).

**Figure 3.**
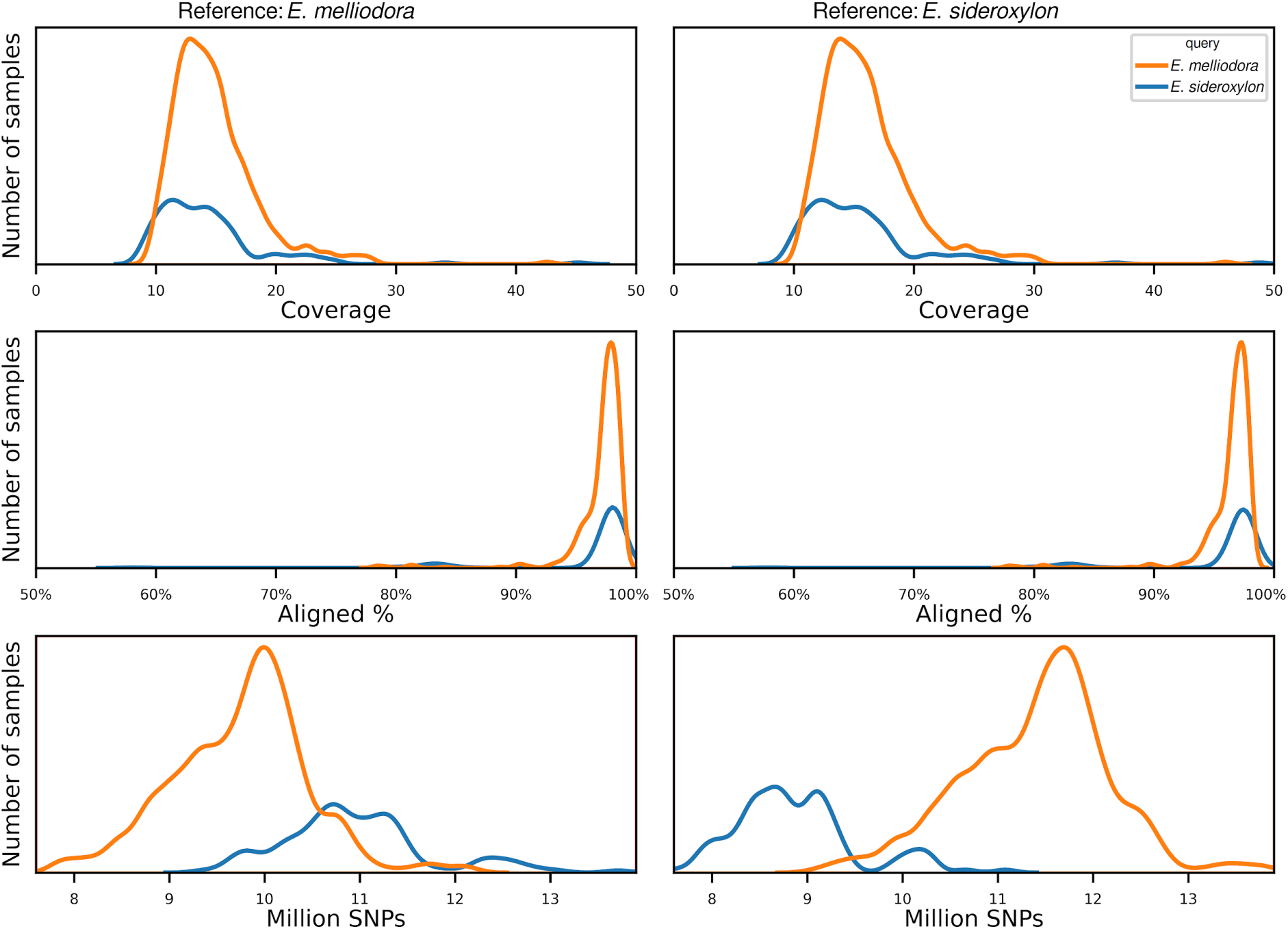
Sample coverage, alignment, and SNP distributions. Left figures use *E. melliodora* as the reference, showing the per sample histogram of sample coverage, percent of reads successfully aligned to reference, and the number of SNPs detected. Right figures are identical, except use *E. sideroxylon* as the reference.

**Figure 4.**
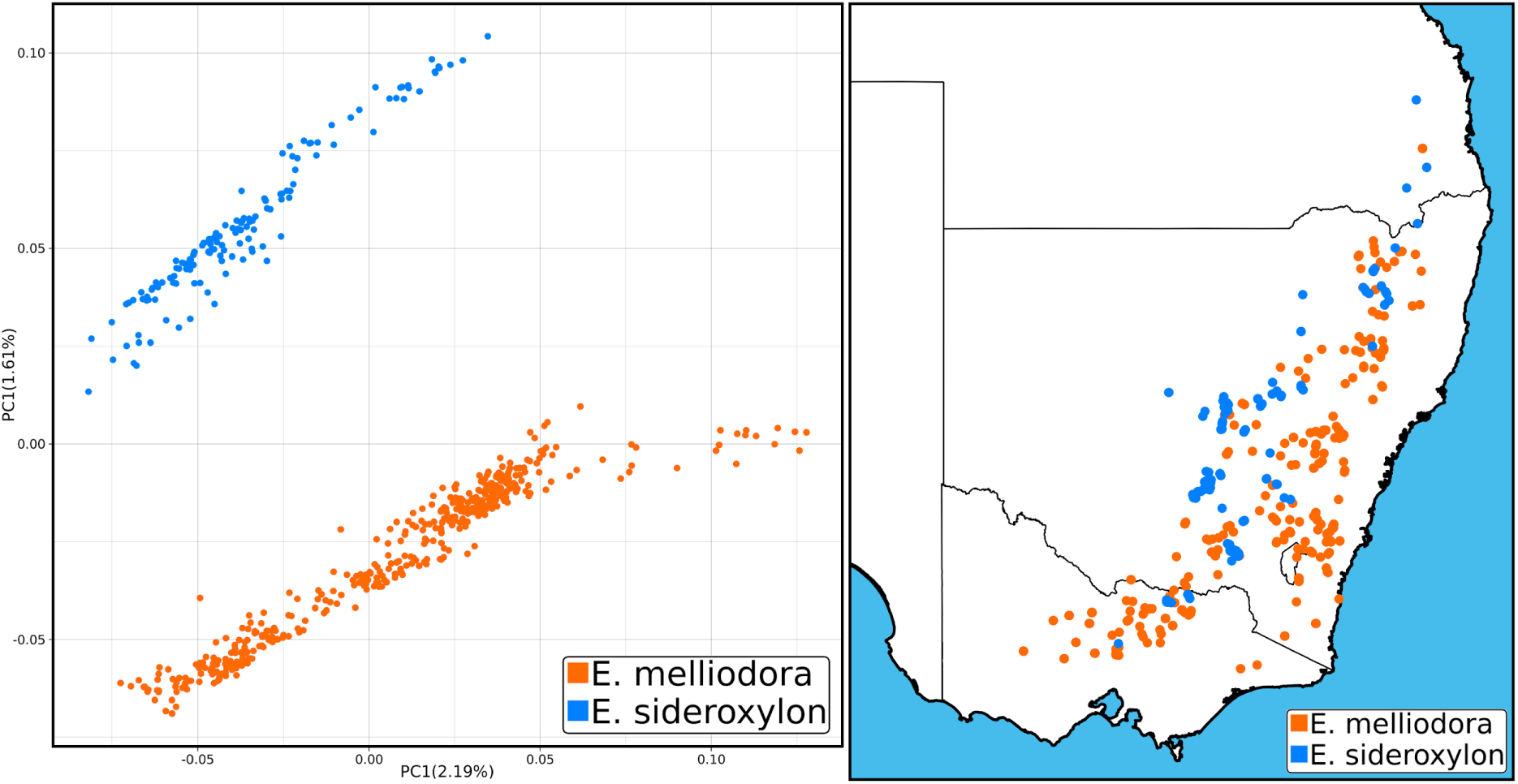
Principal component analysis and sample distribution. Left PCA plot uses *E. melliodora* as the reference genome following the removal of mislabelled, hybrid, and outlier samples. Right map shows the spatial distribution of samples across south eastern Australia. For PCA using *E. sideroxylon* as the reference see Supplementary Figure S5.

**Table 3.** Short-read sequencing, alignment, SNP, and recombination rate estimate statistics.

|  | Reference species | *E. melliodora* |  | *E. sideroxylon* |  |
| --- | --- | --- | --- | --- | --- |
|  | Population species | *E. melliodora* | *E. sideroxylon* | *E. melliodora* | *E. sideroxylon* |
|  | All samples | 459 | 154 | 459 | 154 |
|  | Filtered samples | 425 | 138 | 425 | 138 |
| Estimated read coverage | Average | 14.90 | 16.03 | 14.82 | 15.52 |
|  | Range | 10.00 - 42.58 | 10.59 - 45.97 | 10.11 - 45.16 | 10.06 - 48.76 |
| Read alignment | Average | 97.06% | 96.43% | 96.32% | 95.81% |
|  | Range | 78.40% - 98.80% | 78.70% - 98.56% | 77.91% - 98.10% | 78.38% - 98.02% |
| SNPs (million) | Average | 9.74 | 10.93 | 11.36 | 8.88 |
|  | Range | 6.77 - 13.50 | 8.30 - 13.80 | 7.07 - 14.80 | 7.61 - 12.05 |
|  | Total | 23.16 | 25.39 | 24.96 | 21.87 |
|  | Grand total | 32.46 |  | 31.28 |  |
| Recombination rate estimates | Genome-wide | 0.050 ± 0.031 | - | 0.049 ± 0.033 | - |
|  | Chromosome average range | 0.049 - 0.052 | - | 0.047 - 0.049 | - |

Principal component analysis identified 15 samples that were mislabelled, hybrid, and outlier samples, which were removed (supplementary Figures S4). After removal of these samples the PCA showed two distinct species groups, Figure 4. Within the combined *E. melliodora* dataset, 32.45 million sites, or 5.20% of the genome, were found to be variable. 49.61% of these SNPs were found segregating within both species, 21.76% were private to *E. melliodora,* and as expected a larger proportion, 28.63%, were private to the non-reference species *E. sideroxylon*. Within the combined *E.sideroxylon* SNP dataset we observed the same pattern; 31.28 million SNPs (5.38% of the genome) were found, of which 49.68% segregated within both species, and 20.24% were private to *E. sideroxylon*, while a larger proportion, 30.08%, was found within the non-reference species (Table 3).

### Structural variation genotyping

Interspecies SVs identified between *E. melliodora* and *E. sideroxylon* may be categorised as SD, SP, or SSP. Structural divergences are any event fixed within one species and absent from the other. Structural polymorphisms are any event fixed or absent in one species and polymorphic in the other. Shared structural polymorphisms are SVs that are polymorphic in both populations, Figure 1. Genotyping an SV as SD, SP, or SSP requires examination within both species. While symmetric rearrangements, such as inversions and translocations, can be directly genotyped in both populations, duplications pose challenges due to their asymmetry. Although converting duplications into insertions for short-read genotyping is possible, accurately placing them within the opposite genome is difficult and may result in false negative genotypes. Additionally, genotyping unaligned regions as insertions or deletions introduces uncertainties, especially as they may represent insertions, deletions, or divergent sequences. Short-read alignments with low mapping scores may confound genotyping of unaligned regions (Alser et al., 2021; Valiente-Mullor et al., 2021). Hence, we approach unaligned regions with caution, refrain from categorising duplications as SD, SP, or SSP, and focus our analysis on inversions and translocations for more reliable results. All analyses are performed per allele (2 x population size), not per sample.

Genotyping SVs with short-read alignments resulted in the successful genotyping of 81.11% and 79.46% of SVs in *E. melliodora* and *E. sideroxylon*, respectively (Figure 5). The majority of SVs were found to be fixed (60.65% - 85.10%) or polymorphic (14.84% - 38.57%), with the remaining small proportion (0% - 1.45%) being private to the reference or assembly/scaffolding artefacts. To categorise symmetric interspecies SVs as SD, SP, or SSP we combined the status of fixed inversions (*E. melliodora*: 130; *E. sideroxylon*: 174), polymorphic inversions (*E. melliodora*: 66; *E. sideroxylon*: 37), fixed translocations (*E. melliodora*: 5,652; *E. sideroxylon*: 6,634), polymorphic translocations (*E. melliodora*: 3,288; *E. sideroxylon*: 2,117) across both species, Figure 6. The analysis revealed that the majority of inversions and translocations were either fixed in both species or not successfully genotyped in both species, representing SVs private to the reference genome or assembly/scaffolding artefacts. The remaining proportion consisted of SPs (inversions: 25.98%, translocations: 24.80%) or SSPs (inversions: 7.79%, translocations: 8.81%).

**Figure 5.**
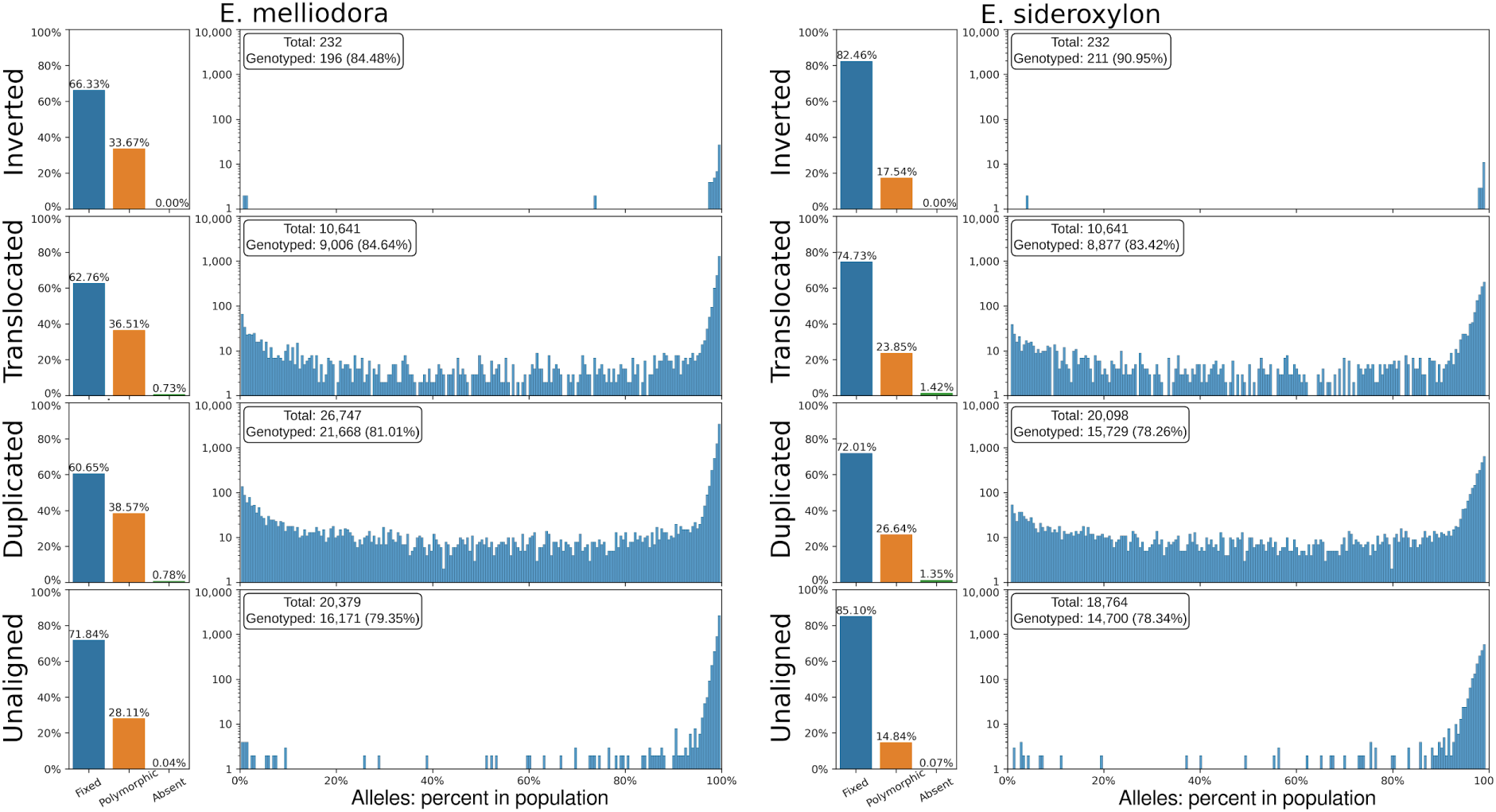
Interspecies SVs and unaligned region frequencies within *E. melliodora* and *E. sideroxylon*.

**Figure 6.**
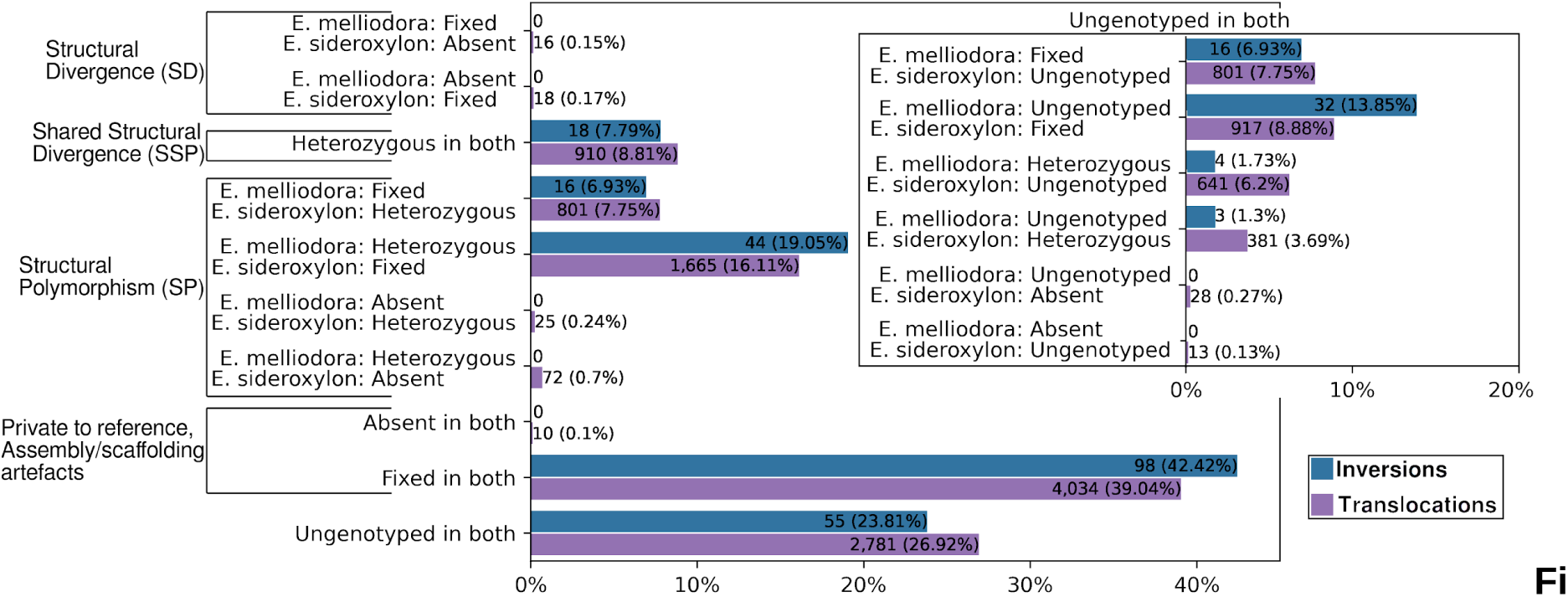
Categorisation of interspecies inversions and translocations as SD, SP, and SPP.

Examination of polymorphic SVs revealed a bimodal distribution of alleles containing the SV, Figure 5. Polymorphic SVs were either very frequently (> 90%) genotyped or very infrequently (< 10%) genotyped within the two species. However, while bimodally distributed, the very frequent SV peak was found to be much higher than the very infrequent.

### Structural variation linkage

Linked variations are those that co-occur more often than would be expected by random chance. Structural variations may be linked by physical proximity, drift or evolution. Evolutionarily linked SVs are likely to contribute to an individual’s survivability and be required for gamete viability and/or the offspring’s adaptive potential. To find evidence of SV linkage we measured correlations among all inversions and translocations for all individuals within both species. For efficient analysis, inversions and duplications were grouped by type (SD, SP, and SSP). Visual inspection of the resulting correlation heatmaps shows several SVs are linked across all categories, Figure 7. To examine the potential role of physical proximity on SV linkage, we examined the distance between correlated SV pairs. 86.64% of SV pairs were found on different chromosomes. When on the same chromosome, SVs were at least 51 Kbp separated.

**Figure 7.**
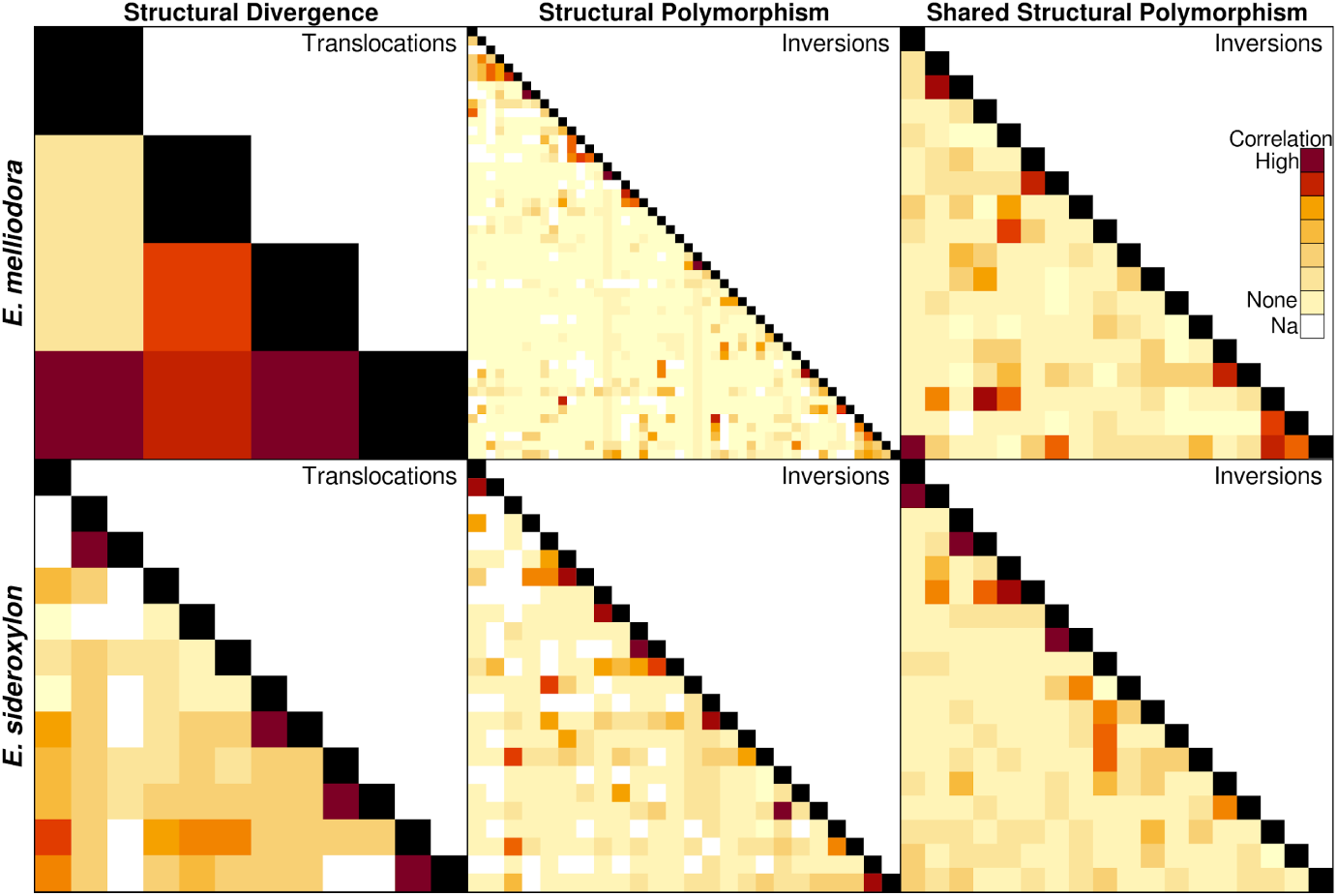
Correlation of SVs between samples. A positive correlation between SV implies that SVs exhibit a non-random association and suggests that these variants tend to co-occur within the population. Categories of SV not present were either empty, as in the case of inversion SD, or contained too many SV to visualise clearly, as in the case of translocation SP and translocation SSP. Undefined correlations, resulting from the failure of short-read to resolve presence/absence of SVs, were removed.

### Shared Structural Polymorphisms COG terms

As SSPs are likely ancestral SVs that have survived drift, underdominant selection, and lineage divergence, they may contain genes of adaptive or other evolutionarily significant value. Examination of SSPs identified 281 gene candidates (*E. melliodora*: 145 and *E. sideroxylon*: 136), of which 247 (87%; *E. melliodora*: 125 and *E. sideroxylon*: 122) were functionally annotated into eggNOG orthogroups and grouped into COG (Clusters of Orthologous Groups; Galperin et al., 2020) categories, Figure 8. Similarly, all genes were functionally annotated and placed within COG categories. Comparing all genes to SSP genes indicates that SSP genes have an increased association with DNA replication, DNA recombination, DNA repair, post-translational modification, protein turnover, chaperones, signal transduction, intercellular communication, and unexplored aspects of biology. Similarly, genes within SSPs have a decreased association with categories for fundamental cellular functions, such as protein synthesis, defence against pathogens, maintaining cellular integrity, providing structural support, and regulating crucial molecular processes involving amino acids, nucleotides, and coenzymes.

**Figure 8.**
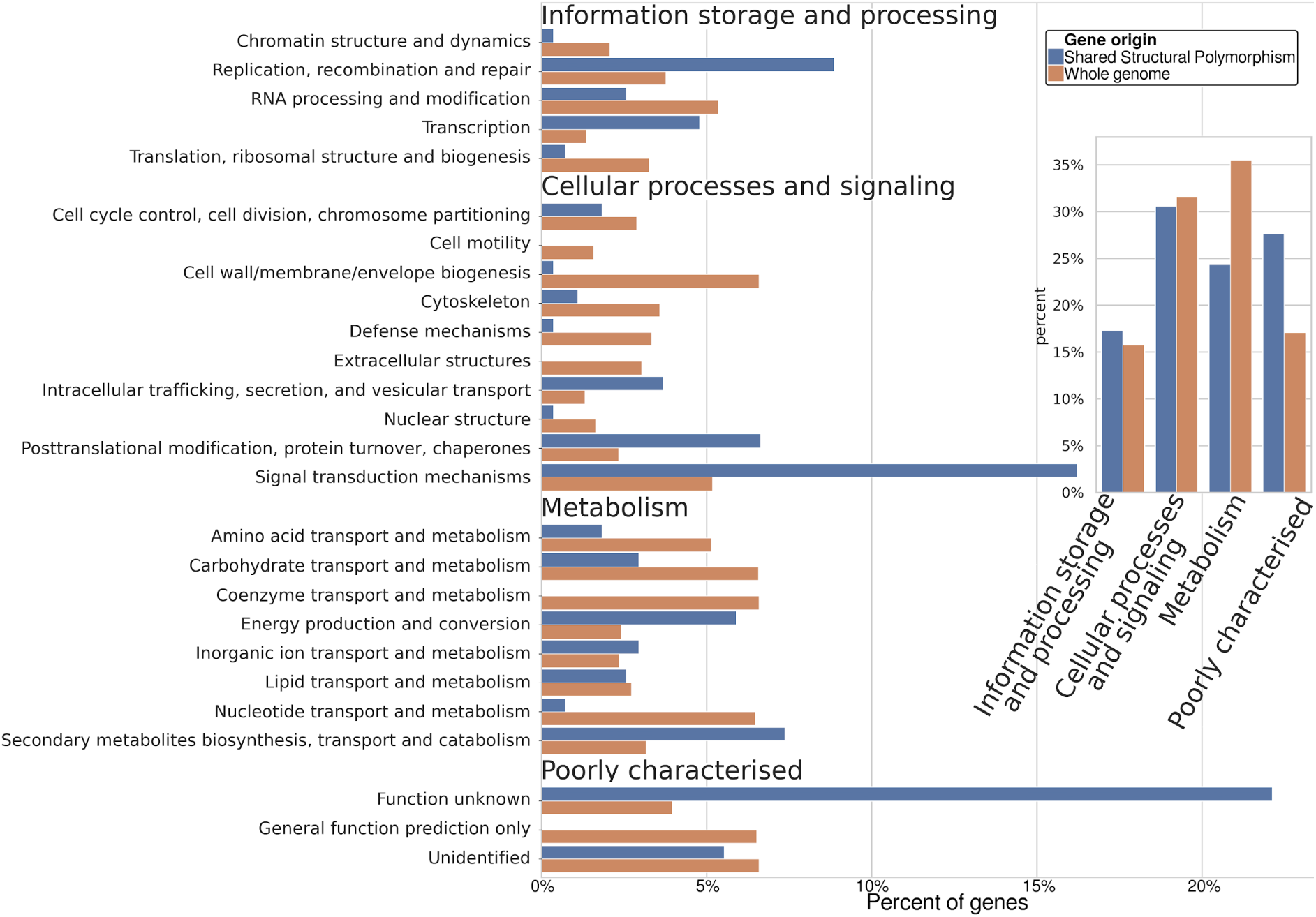
Clusters of Orthologous Groups (COG) terms for all genes and genes found within SSPs.

### Effect of Synteny, rearranged, unaligned, and genes on recombination rates

After annotating SVs in both species and genotyping their frequencies, we calculated ρ across the reference genomes. As low-frequency SVs are unlikely to have a detectable effect on ρ, we considered only fixed SVs and excluded events shorter than 2 Kbp, as ρ was calculated within 1 Kbp windows. We also assessed the impact of genes and transposons larger than 2 Kbp on ρ. Prior to ρ calculations, we phased SNPs, initially achieving 20.56% linkage within haplotype blocks using read alignments, and subsequently completing phasing with a HMM-based approach. After separation of SNPs into parental haplotypes, we found that *E. sideroxylon* consistently exhibited higher and more variable ρ compared to *E. melliodora*. Chromosome-specific recombination rates displayed notable variability without discernible patterns, Table 3, Supplementary Table S2, and Supplementary Figures S6 and S7.

An initial ANOVA assessment indicated differences in ρ for our different categories of genome regions, for both species (p-value; *E. melliodora*: 8.35×10^-276^ and *E. sideroxylon*: 1.85×10^-272^). To determine if any region type/s were contributing to differences in ρ, we performed Tukey’s test, adjusting p-values to account for the total species error rate. Tukey’s test for *E. melliodora* revealed that, in comparison to syntenic regions, average ρ was higher for genes, transposons, inversions, and duplications, Figure 9A. However, statistically significant differences were observed only for genes, transposons, and duplications. Notably, genes followed by transposons exhibited significantly higher ρ than all other types of regions, while duplications showed higher values than translocations and unaligned regions. Inversions exhibited a wider confidence interval (CI) due to their lower number of events. A similar pattern was observed by Tukey’s test for *E. sideroxylon*. While genome-wide statistical observations of ρ were unrevealing, many SVs were observed having ρ less than the mean syntenic, Figure 9C and 9D. For detailed significance testing results refer to Supplementary Table S3.

**Figure 9.**
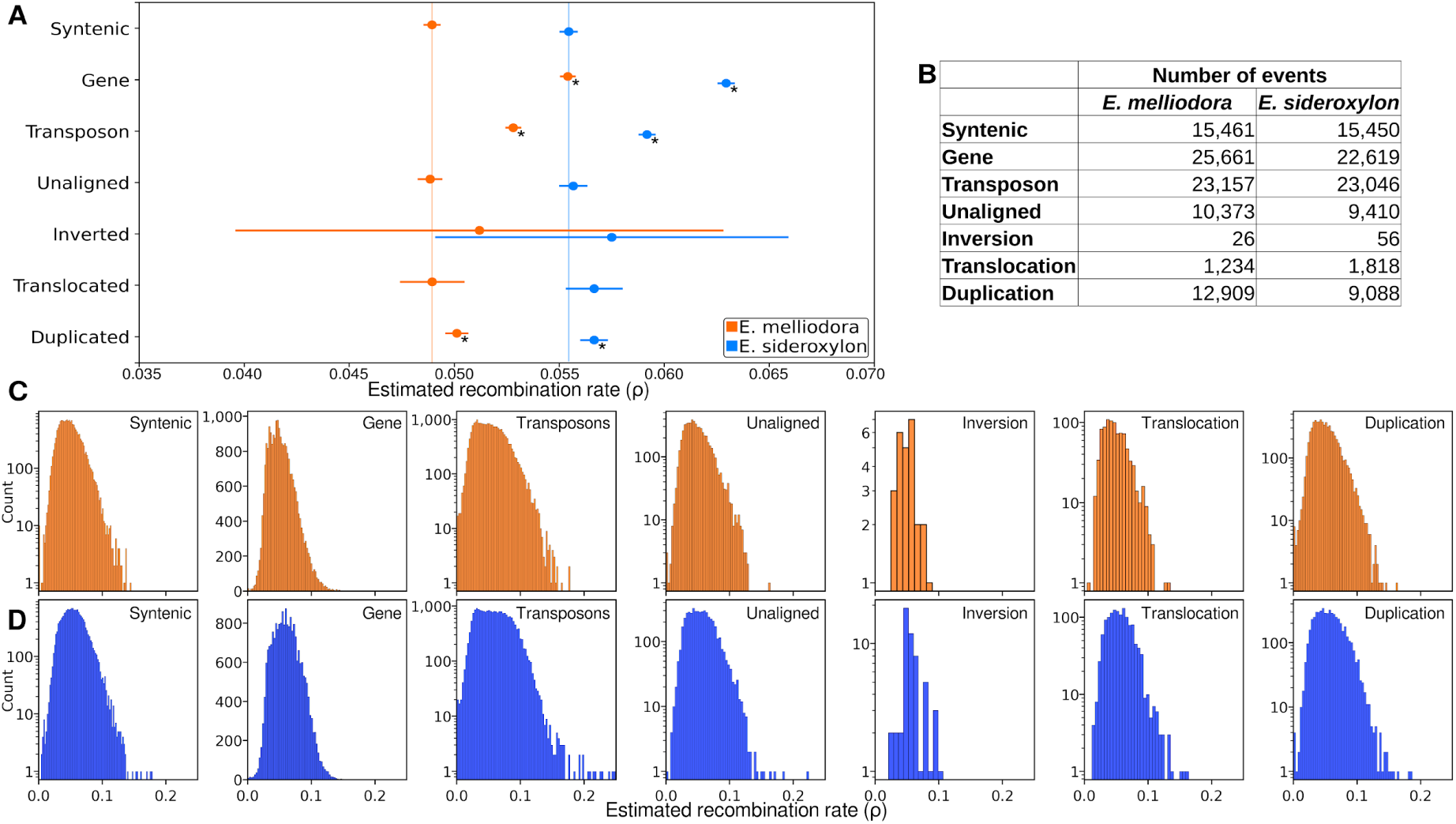
Tukey’s test for estimated recombination rates of fixed SVs, unaligned regions, genes, and transposons. A) Shows mean and 95% confidence interval for all events. Vertical lines show average ρ for syntenic regions. * indicates region types that are significantly different from syntenic regions (P ≤ 0.05). B) The number of events included in the analysis. C) Estimated recombination rate distribution for *E. melliodora*. D) Estimated recombination rate distribution for *E. sideroxylon*.

### Effect of Synteny, rearranged, unaligned, and genes on Fixation index (*F_ST_*)

To measure the amount of shared genetic diversity that exists between *E. melliodora* and *E. sideroxylon*, we combined SNPs for both populations under each reference and calculated the fixation index (*F_ST_*). The fixation index, calculated per SNP, scores the amount of genetic differentiation between populations or species and ranges from 0 to 1, where 0 indicates no difference in allele frequencies and 1 indicates a fixed difference. In real world usage, per SNP *F_ST_* values are typically far below one, even in the case of isolated populations and should be interpreted relative to the study (Kitada et al., 2021). Here we use them to quantify how similar, or dissimilar, all region types are between *E. melliodora* and *E. sideroxylon*.

As per our examination of ρ, we calculated the average *F_ST_* for all fixed SVs, and genes and transposons greater than 2 Kbp in length and performed Tukey’s test, Figure 10A. Syntenic regions were used as the reference point to evaluate the extent of genetic differentiation of SVs. Using *E. melliodora* as the reference, all region types had significantly less divergence between species except genes and inversions. Genes had significantly more divergence and inversions were sparse and as such had a wide confidence interval. A similar pattern was observed for *E. sideroxylon*. While genome-wide statistical observations of F_ST_ were unrevealing, many SVs were observed having F_ST_ less than the mean syntenic, Figure 10C and 10D. Examination of F_ST_ histograms for all event types showed a left shifted Poisson distribution, with many events having low F_ST_ scores. For detailed significance testing results refer to Supplementary Table S4.

**Figure 10.**
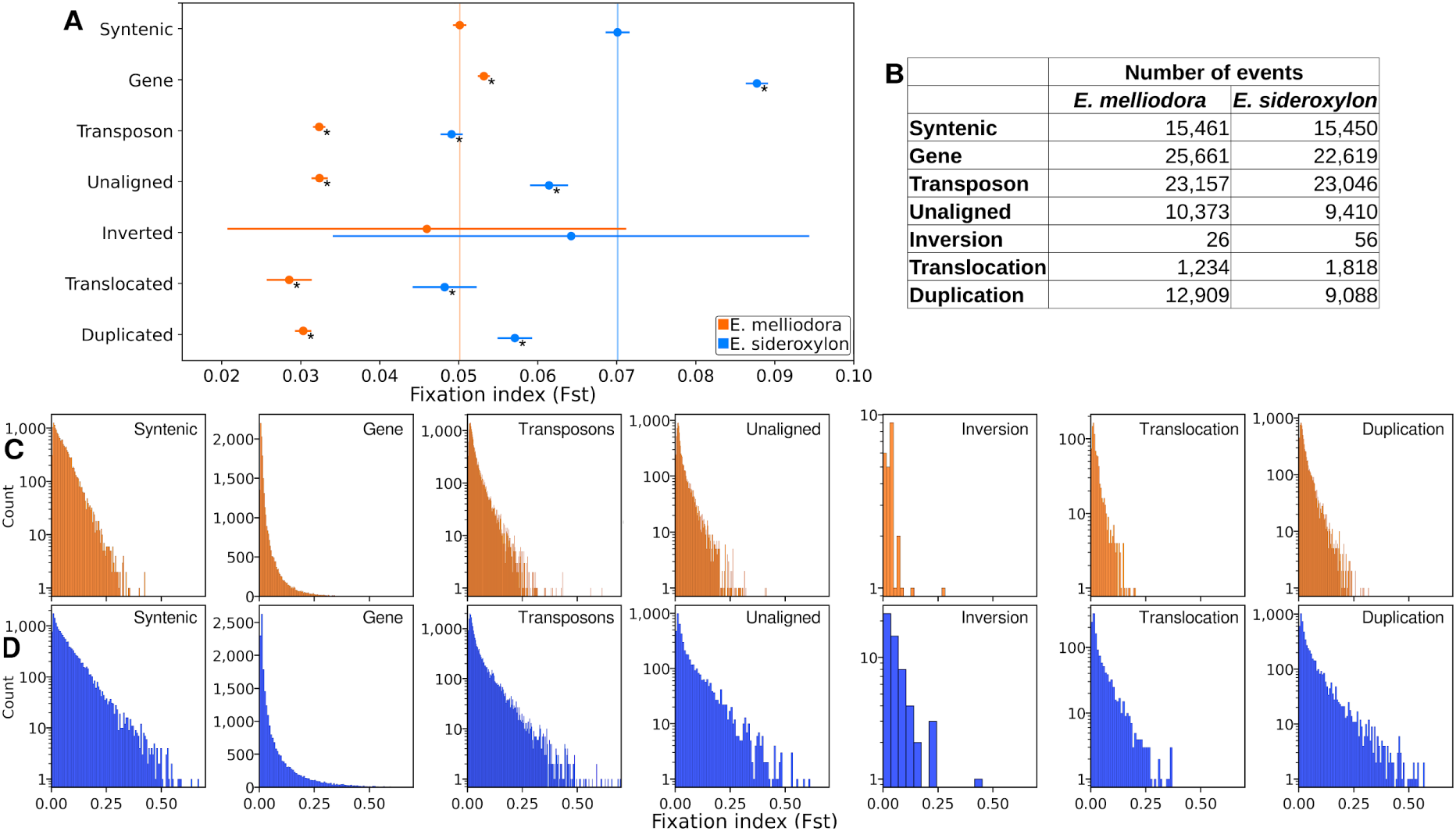
Tukey’s test for Fixation index of fixed SVs, unaligned regions, genes, and transposons. A) Shows average F_ST_ and 95% confidence intervals calculated from average F_ST_ values for all regions. For each reference genome, SNPs from both species were combined and F_ST_ calculated. Vertical lines show average ρ for syntenic regions. * indicates region types that are significantly different from syntenic regions (P ≤ 0.05). B) The number of events included in the analysis. C) Fixation index distribution for *E. melliodora*. D) Fixation index distribution for *E. sideroxylon*.

### Effect of Synteny, rearranged, unaligned, and genes on SNPs

SNP density can significantly impact the precision and resolution of both ρ and F_ST_ (Akey et al., 2002; Bhatia et al., 2013; Chan et al., 2012). Higher SNP density enables finer-scale mapping of recombination events and more accurate population differentiation measurements, while lower SNP density gives coarser results with reduced precision. Due to inconclusive results in both ρ and F_ST_ analyses, we examined SNP densities of SVs, genes, and TEs.

As per our ρ and F_ST_ analyses, we used Tukey’s test and histograms to examine the differences in SNP densities for all fixed SVs, and genes and transposons greater than 2 Kbp in length, Figures 11A, 11C, and 11D. For detailed significance testing results refer to Supplementary Table S5. Reassuring to our SV annotation method, unaligned regions were the most diverged region type, containing the largest number of SNPs. Similarly reassuring for our annotation method, genes were the least diverged, containing the fewest SNPs. No significant correlations between the number of SNPs and ρ were observed. Notably, genes, transposons and duplications had high ρ, while only transposons had a high SNP density. Conversely, unaligned and translocated regions had low ρ, while only translocation had few SNPs. Similarly, no distinct correlations between SNPs and F_ST_ values were observed. Genes, despite having few SNPs, contained high F_ST_ values, whereas unaligned regions, with many SNPs, displayed low F_ST_ values. Translocated regions, with an intermediate number of SNPs, also exhibited low F_ST_ values. Although SNP densities contribute to the complex pattern of genomic differentiation, they showed no clear association with ρ and F_ST_ calculations.

**Figure 11.**
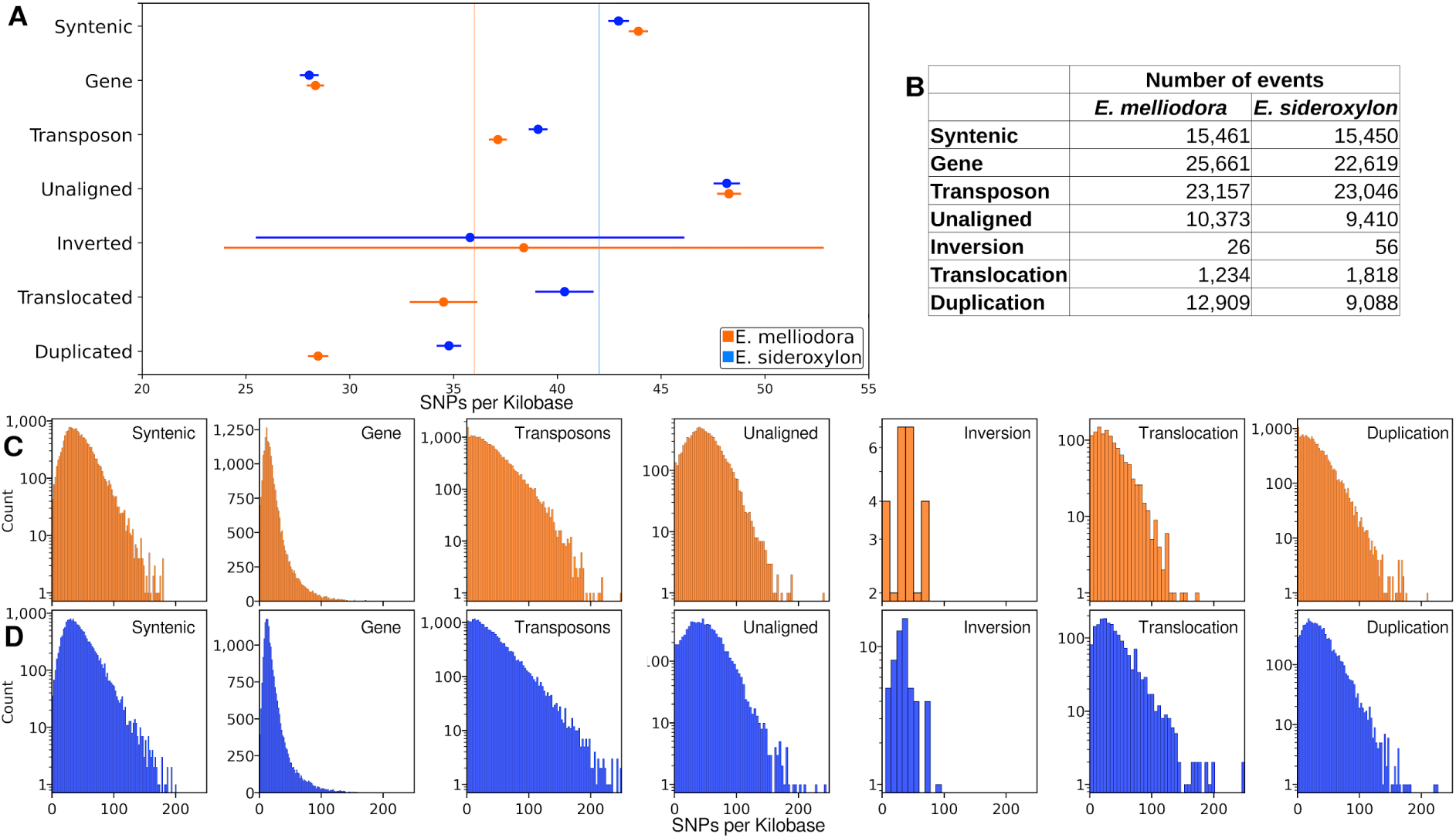
Tukey’s test for SNP density of fixed SVs, unaligned regions, genes, and transposons. A) Shows mean and 95% confidence interval for all events. Vertical lines show average ρ for syntenic regions. B) The number of events included in the analysis. C) SNP density distribution for *E. melliodora*. D) SNP density distribution for *E. sideroxylon*.

## Discussion

Structural variants are a major form of genomic variation, affecting more nucleotides than SNPs (Escaramís et al., 2015). Despite their prominence, the functional and evolutionary impacts of SV’s remain poorly understood (Chain & Feulner, 2014; Ho et al., 2020; Yan et al., 2021). To date, the majority of population-scale SV studies have focused on within population SV discovery and association with environments or phenotypes (Gui et al., 2022; Hufford et al., 2021). Several studies have also directly examined SV and their contribution to functional changes (Ishikawa et al., 2019; Zhao et al., 2020). Here we genotyped interspecies SVs and described their frequencies within and among both species. Of particular novelty is our comparison of translocations and inversions, symmetric SVs that may be present within one or both species and at different frequencies. Between our recently diverged *Eucalyptus* species pair, our results demonstrate that SVs contribute to genome divergence, intra-species genetic diversity, and shared genetic diversity. Potentially of great interest are shared structural polymorphisms (SSP); these large mutations predate lineage divergence and remain polymorphic within both species, potentially containing locally adaptive or otherwise important genes and allele combinations. Additionally, examination of average ρ and *F_ST_* within fixed SVs demonstrates the variable effects of these genetic variations on genome differentiation and recombination.

Genetic mutations promote and reinforce lineage divergence, are the genetic basis of reproductive isolation, and are essential to the process of speciation. Structural mutations, by affecting recombination, phenotypes, or altering/removing/sub-functionalising genes, are of particular importance to speciation processes (Zhang et al., 2021). Barrier complexity and asymmetry are underappreciated components of reproductive isolation. Barrier complexity involves the combinatorial interplay of genetic barriers that collectively reduce reproductive success between individuals (Shang et al., 2020). Successful offspring are survivors of genetic combinations, possessing genomes sufficiently free from barrier loci to allow reproduction to occur. Barrier asymmetry refers to the relative effectiveness of reproductive barriers between two groups, resulting in different hybridisation success rates (Christie et al., 2022). *Eucalyptus melliodora* and *E. sideroxylon* are known to hybridise, and successful hybridisation likely results from the complex interplay between the numerous SD, SP, and SPP that come together in a particular hybrid. Evidence of linked SDs, SPs, and SPPs was observed within both *Eucalyptus* species. These linked SV combinations may be required for reproductive success or there could be some other fitness consequence that is maintaining selection for that linked state. Barrier SVs potentially exhibit a higher degree of reproductive isolation compared to non-SV regions, increasing genetic differentiation within these loci (Berg et al., 2016; Huang et al., 2020; Lucek et al., 2019). However, variation in *F_ST_* did not provide sufficient evidence on average to support this conclusion, possibly due to the recent divergence of our species and the importance of only a few key interacting loci.

Similar to reproductive isolation, understanding how all types of genetic mutations contribute to the creation and maintenance of genetic diversity is crucial to understanding how organisms improve fitness and adapt to their changing environments (Gregory, 2009; Loewe & Hill, 2010; Pokrovac & Pezer, 2022). Inversions and translocations aid in adaptive evolution by fixing allele combinations, duplications contribute to the development of new genes, and insertions and deletions, often described as PAVs, modify gene expression and gene content (De Oliveira et al., 2020; Mérot et al., 2020; Wellenreuther et al., 2019). A substantial number of inversions and translocations were successfully genotyped within both species. The majority of inversions and translocations were SP or SSP, making them candidates for exploring adaptive genes and alleles. Of particular note are SSP inversions and translocations which showed evidence of gene enrichment in potentially adaptive genes.

Duplications are known to be highly common and an important source of evolutionary novelty (Cohen et al., 2023; Hanada et al., 2008), and were the most common type of SV in our analysis. Most duplications were found to be fixed, with the remainder being almost entirely polymorphic. Given their asymmetry, duplications were genotyped only within their respective host genomes, resulting in an inability to categorise them as SD, SP, or SSP. Nonetheless, duplications successfully genotyped in our study are potential candidates for adaptive loci, likely having withstood the influences of genetic drift and purifying selection. Predicting the adaptive effects of unaligned regions presents a significant challenge, given their potential to encompass insertions, deletions, or highly divergent sequences. When unaligned regions result from highly divergent sequences, short reads will align poorly, confounding genotyping (Alser et al., 2021; Valiente-Mullor et al., 2021). Genotyped as deletions, the majority of unaligned regions were fixed, and the remainder highly frequent. Fixed unaligned regions are possibly highly diverged regions or a deletion in the opposite species genome. When unaligned regions are polymorphic could represent insertions within the host species genome or deletions within the opposite species genome. These difficult to interpret regions may be PAVs, adaptive loci, or selectively neutral or deleterious loci undergoing potential purifying selection. Further investigations are essential to uncover their precise roles and implications.

It is now clear that SVs are of great evolutionary importance and must be considered when studying genetic diversity and genome evolution (Wellenreuther et al., 2019). To better evaluate the impact of SVs on evolution a combination of interspecies and intraspecies studies are crucial. While structural polymorphisms may be reproductive barriers or adaptive loci, they could also be neutral or deleterious, especially as these species separated very recently. Given that SVs are rarely conserved (i.e., typically purged over short time scales) (Inoue et al., 2015; Naseeb et al., 2017), and many of the SV examined here were genotyped at high frequencies, there is potential for common SVs to be investigated for functional associations with traits or environments, thus warranting future scrutiny regarding their contribution to adaptive evolution. Future studies are needed to test whether these SVs contribute to adaptive evolution. To assess their potential role as barrier loci, breeding experiments could be employed. A problem encountered here was the number of SVs within individuals that could not be genotyped. Many statistical tests require all samples to be genotyped for all genetic variants, employing imputation to fill in missing genotypes. However, all current imputation processes are designed for SNPs captured within haplotype blocks. Statistical association programs that can incorporate SVs are needed. With the decreasing cost and increasing accuracy of long-read sequencing, particularly Oxford Nanopore (Ferguson et al., 2022), future studies could utilise high-throughput long-read sequencing to overcome the limitations of short-read SV genotyping. However advances in analysis software are still a limiting constraint for fully understanding the contribution of SVs to adaptive evolution and speciation.

## Data access

Sequencing data and reference genomes generated in this project are publicly available on the Sequence Read Archive (SRA) and NCBI genome repository under BioProject PRJNA509734 and PRJNA578806. Gene predictions and repeat annotations have been deposited in FigShare and are available at: https://figshare.com/projects/Exploring_polymorphic_interspecies_structural_variants_in_Eucalyptus_Unravelling_Their_Role_in_Reproductive_Isolation_and_Adaptive_Divergence_/183577. All analysis scripts created and used by this project have been deposited within our github repository: https://github.com/fergsc/Polymorphic-interspecies-SVs. Sample metadata including GPS locations are also available at our FigShare.

## Competing interest statement

The authors declare that they have no competing interests.

## Funding

This work was supported by the Australian Research Council (CE140100008; DP150103591; DE190100326) and an Australian Government Research Training Program scholarship.

## Authors’ contributions

Scott Ferguson led the project and ran all the analysis. The project was conceived and designed by all authors. Scott Ferguson wrote the first manuscript draft. All authors contributed to writing and review of the final manuscript.

## Acknowledgements

This research was undertaken with the assistance of resources from the National Computational Infrastructure (NCI Australia), an NCRIS enabled capability supported by the Australian Government.

We would like to thank the Australian National Botanic Gardens in Canberra, Australia for providing plant samples and associated metadata for the two reference genomes, *E. melliodora* and *E. sideroxylon*. This research acknowledges the support provided by the Director of National Parks, the park staff of the Australian National Botanic Gardens, and Parks Australia. The views expressed in this document do not necessarily represent the views of the Australian Government.

We thank David Stanley and Cynthia Torkel for their technical assistance in the laboratory and their friendship throughout the times.

## Notes

### Competing Interest Statement

The authors have declared no competing interest.

