## Supplementary for "Exploring polymorphic interspecies structural variants in Eucalyptus: Unravelling Their Role in Reproductive Isolation and Adaptive Divergence"

| | MAPQ >= 30 | MAPQ > 0 |
|---|---|---|
| Sequenced Read Pairs | 151,590,503 | 151,590,503 |
| Normal Paired | 45,342,881 (29.91%) | 45,342,881 (29.91%) |
| Chimeric Paired | 52,358,576 (34.54%) | 52,358,576 (34.54%) |
| Chimeric Ambiguous | 25,426,118 (16.77%) | 25,426,118 (16.77%) |
| Unmapped | 28,462,928 (18.78%) | 28,462,928 (18.78%) |
| Ligation Motif Present | 0 (0.00%) | 0 (0.00%) |
| Alignable (Normal+Chimeric Paired) | 97,701,457 (64.45%) | 97,701,457 (64.45%) |
| Unique Reads | 40,318,503 (26.60%) | 40,318,503 (26.60%) |
| PCR Duplicates | 54,402,352 (35.89%) | 54,402,352 (35.89%) |
| Optical Duplicates | 2,980,602 (1.97%) | 2,980,602 (1.97%) |
| Library Complexity Estimate | 46,306,556 | 46,306,556 |
| Intra-fragment Reads | 0 (0.00% / 0.00%) | 0 (0.00% / 0.00%) |
| Below MAPQ Threshold | 21,810,955 (14.39%) | 14,070,582 (9.28%) |
| Hi-C Contacts | 18,507,548 (12.21%) | 26,247,921 (17.32%) |
| Ligation Motif Present | 0 (0.00% / 0.00%) | 0 (0.00% / 0.00%) |
| 3' Bias (Long Range) | 50% - 50% | 50% - 50% |
| Pair Type %(L-I-O-R) | 25% - 25% - 25% - 25% | 24% - 26% - 26% - 24% |
| Inter-chromosomal | 9,612,532 (6.34%) | 14,833,362 (9.79%) |
| Intra-chromosomal | 8,895,016 (5.87%) | 11,414,559 (7.53%) |
| Short Range (<20Kb) | 6,771,738 (4.47%) | 8,386,180 (5.53%) |
| Long Range (>20Kb) | 2,113,340 (1.39%) | 3,016,215 (1.99%) |

**Supplementary Table S1.** *E. melliodora* Hi-C summary stats, produced by Juicer.

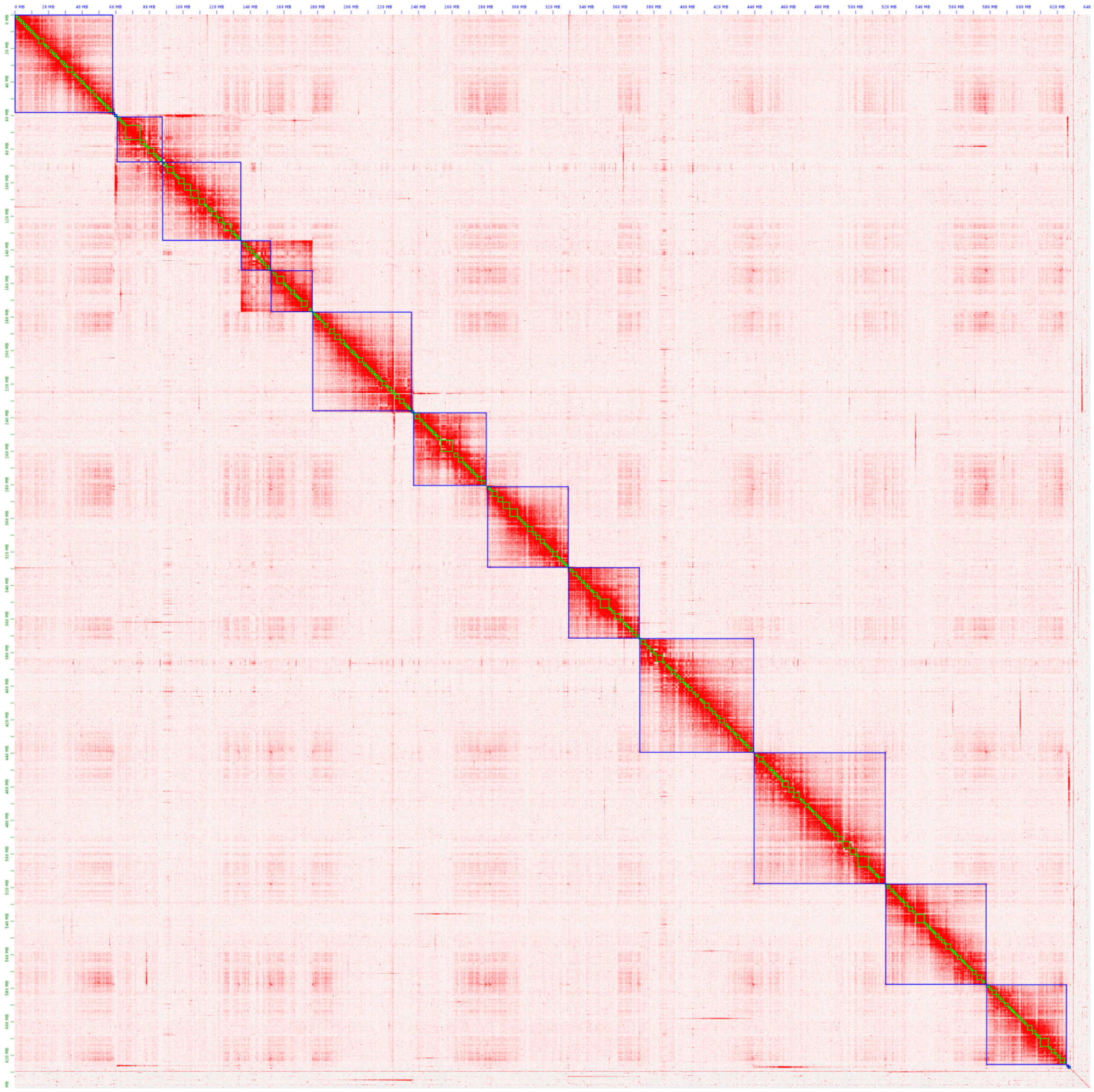

**Supplementary Figure S1.** Hi-C scaffolding of *E. melliodora's* contigs with 3D DNA (parameter: "--editor-repeat-coverage 5, -i 1000). Due to a high repeat content Hi-C read coverage is highly variable, resulting in poor scaffolding. Hi-C contacts are visualised with Juicebox.

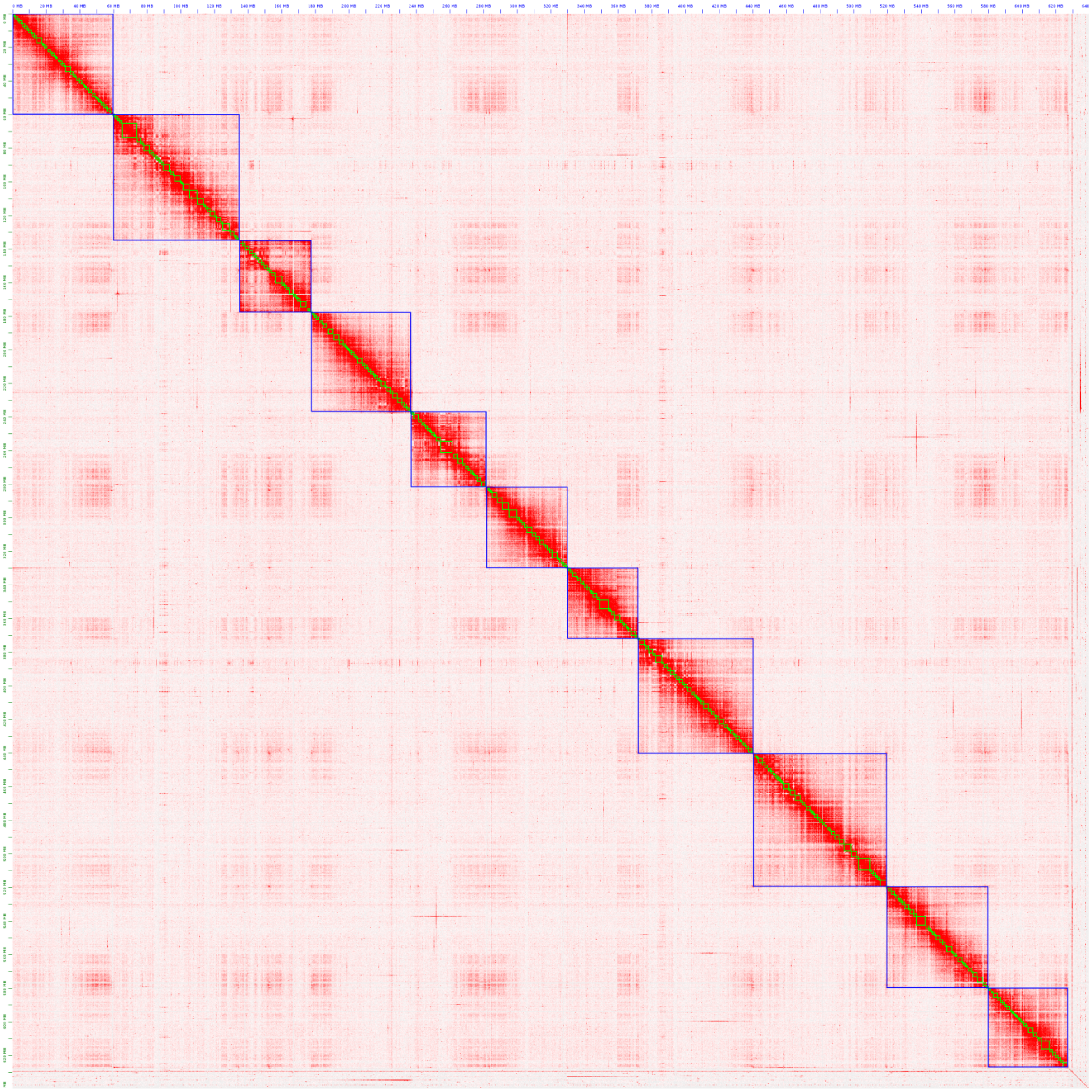

**Supplementary Figure S2.** Manually curated, final, Hi-C contact map of *E. melliodora's* contigs with 3D DNA (parameter: "--editor-repeat-coverage 5, -i 1000). Due to a high repeat content Hi-C read coverage is highly variable, resulting in poor scaffolding. Hi-C contacts are visualised with Juicebox.

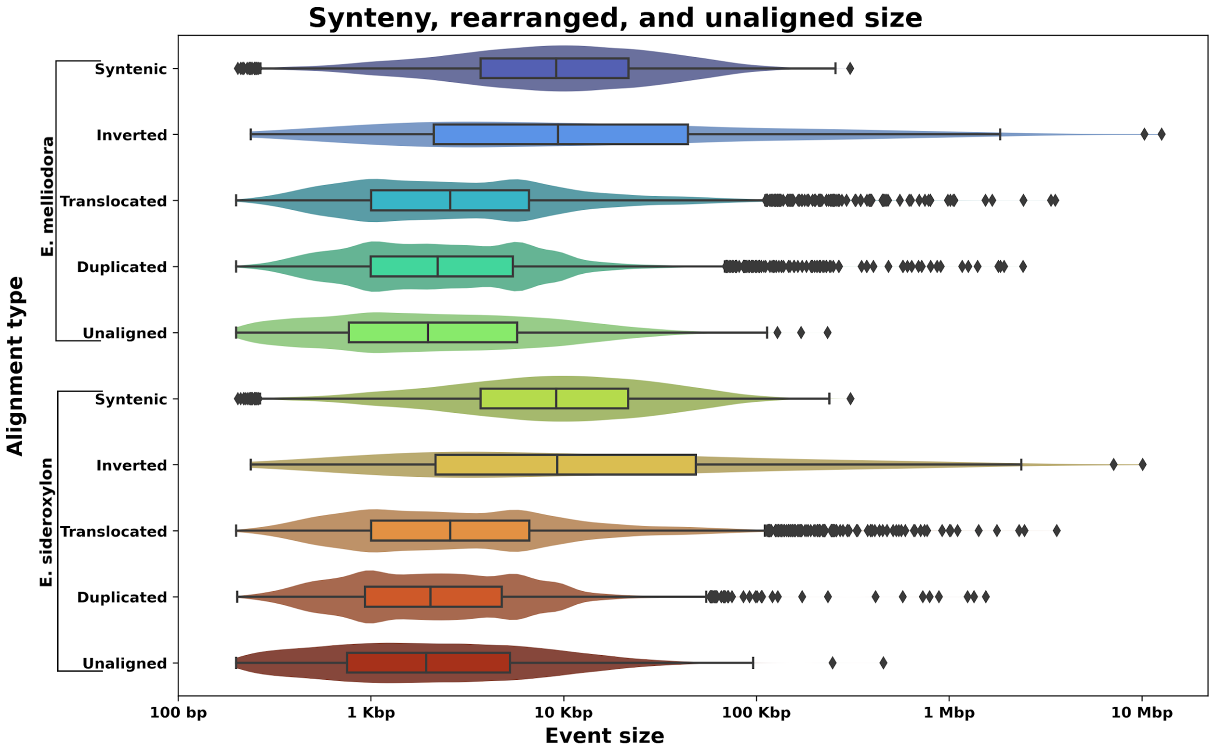


**Supplementary Figure S3.**

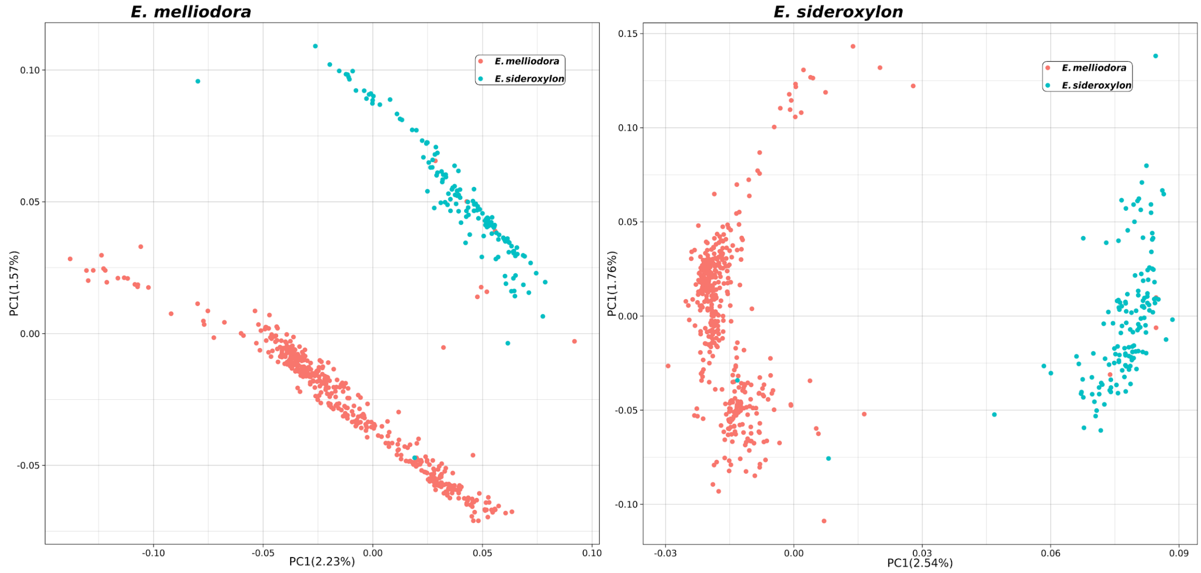


**Supplementary Figure S4. Raw PCA plots.** Left figure uses *E. melliodora* as the reference, the right figure uses *E. sideroxylon* as the reference.

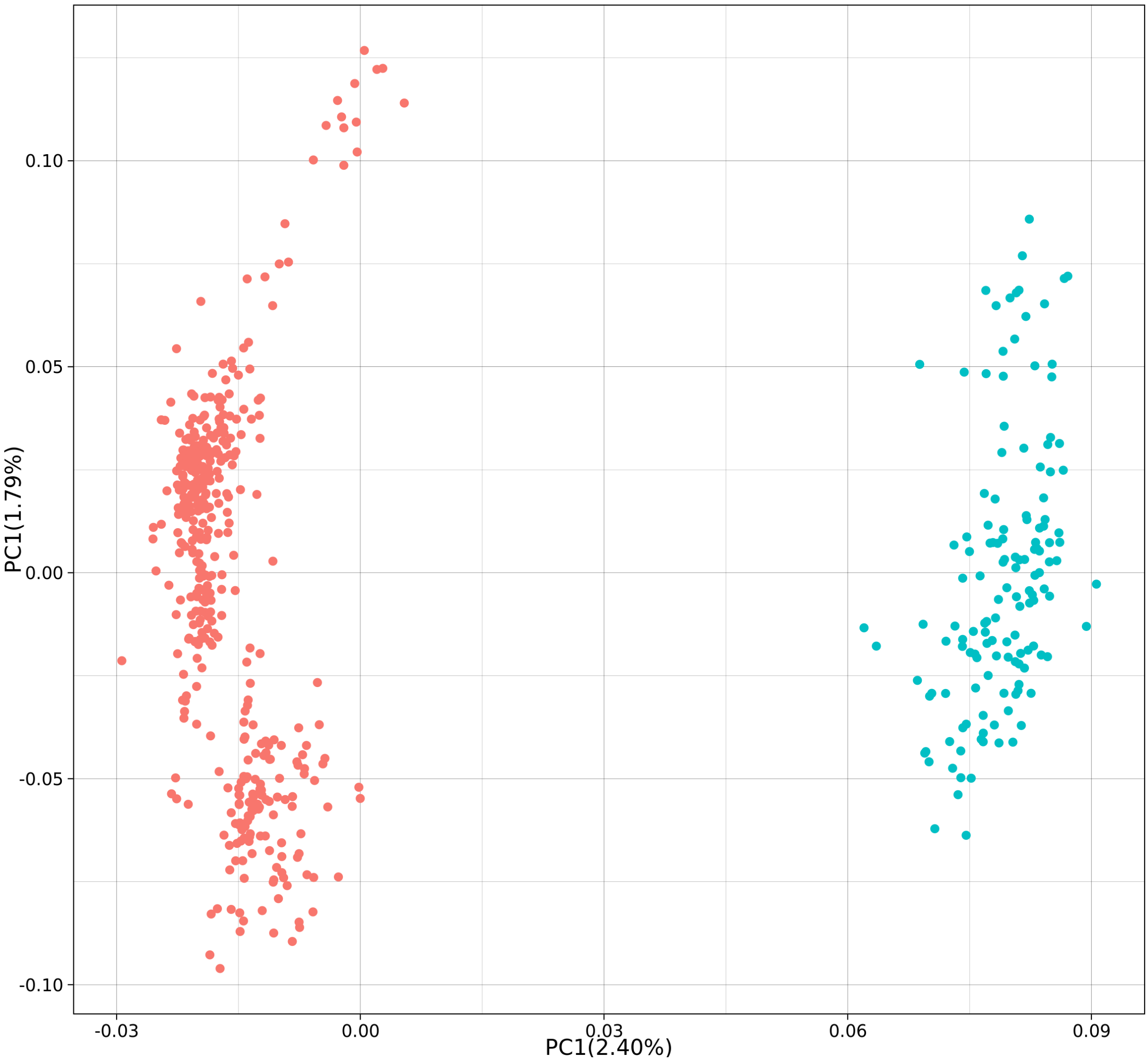


**Supplementary Figure S5. Clean PCA plot, *E. Sideroxylon* as reference.**

| | *E. melliodora* | *E. melliodora* |
|---|---|---|
| **Genome-wide** | 0.05016 ± 0.03074 | 0.04879 ± 0.03188 |
| **Chromosome 1** | 0.04996 ± 0.03134 | 0.04782 ± 0.03180 |
| **Chromosome 2** | 0.05012 ± 0.03069 | 0.04989 ± 0.03180 |
| **Chromosome 3** | 0.04871 ± 0.02954 | 0.04803 ± 0.03136 |
| **Chromosome 4** | 0.04978 ± 0.03159 | 0.04944 ± 0.03360 |
| **Chromosome 5** | 0.04919 ± 0.02971 | 0.04714 ± 0.03009 |
| **Chromosome 6** | 0.05196 ± 0.03126 | 0.04999 ± 0.03327 |
| **Chromosome 7** | 0.04946 ± 0.02985 | 0.04899 ± 0.03079 |
| **Chromosome 8** | 0.05002 ± 0.02996 | 0.04826 ± 0.03074 |
| **Chromosome 9** | 0.05012 ± 0.03186 | 0.04746 ± 0.03142 |
| **Chromosome 10** | 0.05197 ± 0.03194 | 0.05090 ± 0.03439 |
| **Chromosome 11** | 0.05167 ± 0.03208 | 0.04896 ± 0.03282 |
| **Range of average** | 0.04871 - 0.05167 | 0.04714 - 0.04896 |

**Supplementary Table S2.** Recombination rate estimates. Recombination rates were calculated in 1 Kbp windows and averaged across chromosomes. Rates are shown with standard deviation. Chromosomes coloured with darker green have higher average recombination rates.

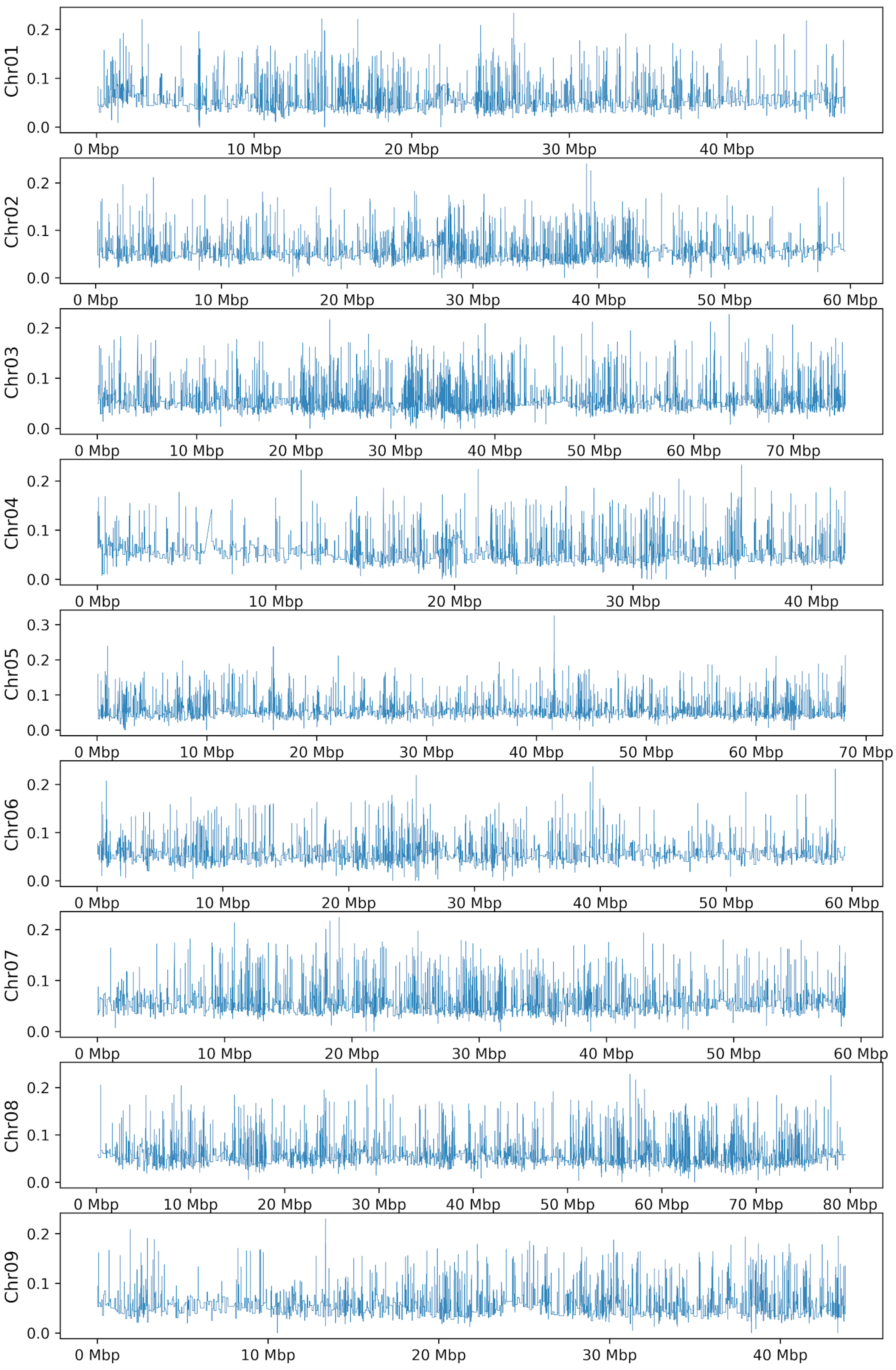
Chr01
0.0
0.1
0.2
0 Mbp
10 Mbp
20 Mbp
30 Mbp
40 Mbp
Chr02
0 Mbp
10 Mbp
20 Mbp
30 Mbp
40 Mbp
50 Mbp
60 Mbp
Chr03
70 Mbp
Chr04
Chr05
0.3
Chr06
Chr07
Chr08
80 Mbp
Chr09

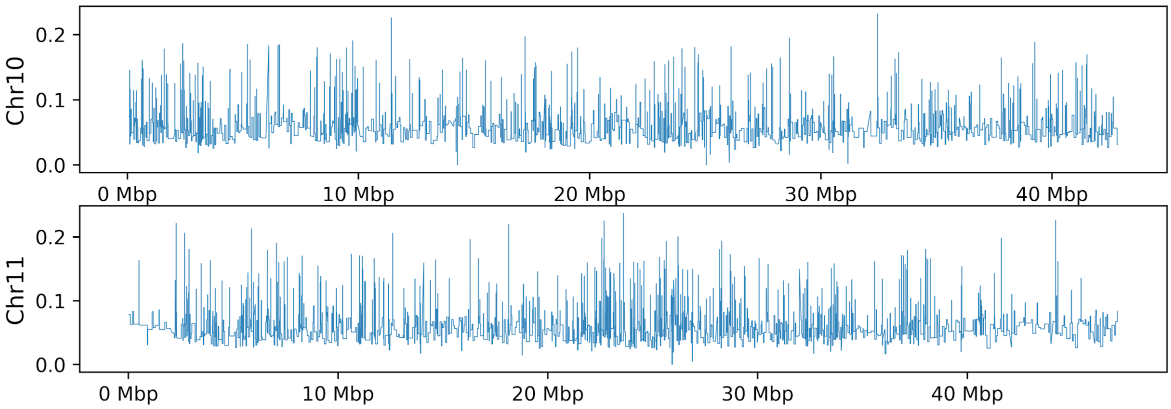


**Supplementary Figure S6.** ***E. melliodora*** **recombination**

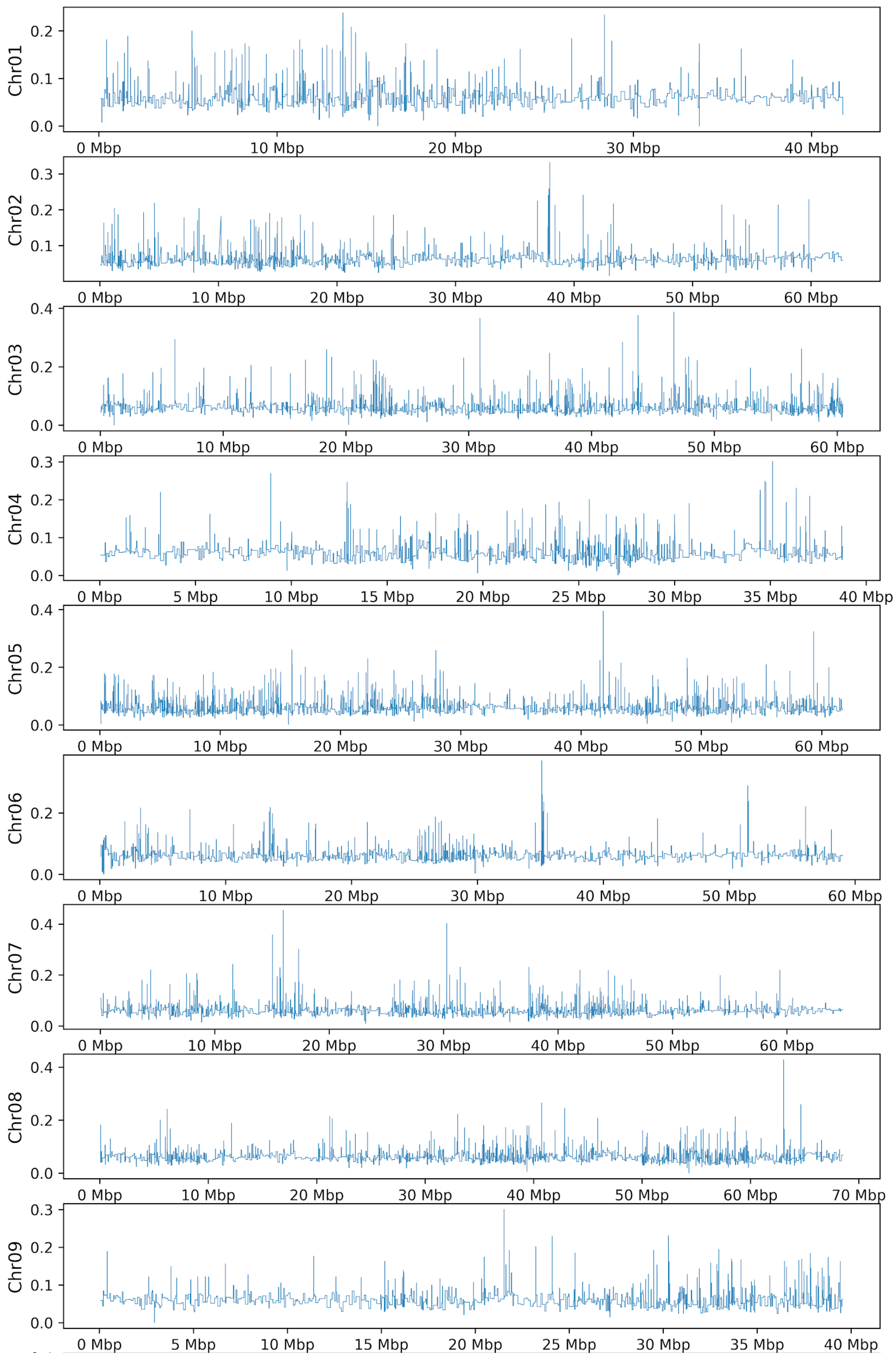

Chr01
0.0
0.1
0.2
0 Mbp
10 Mbp
20 Mbp
30 Mbp
40 Mbp
Chr02
0.1
0.2
0.3
0 Mbp
10 Mbp
20 Mbp
30 Mbp
40 Mbp
50 Mbp
60 Mbp
Chr03
0.0
0.2
0.4
0 Mbp
10 Mbp
20 Mbp
30 Mbp
40 Mbp
50 Mbp
60 Mbp
Chr04
0.0
0.1
0.2
0.3
0 Mbp
5 Mbp
10 Mbp
15 Mbp
20 Mbp
25 Mbp
30 Mbp
35 Mbp
40 Mbp
Chr05
0.0
0.2
0.4
0 Mbp
10 Mbp
20 Mbp
30 Mbp
40 Mbp
50 Mbp
60 Mbp
Chr06
0.0
0.2
0 Mbp
10 Mbp
20 Mbp
30 Mbp
40 Mbp
50 Mbp
60 Mbp
Chr07
0.0
0.2
0.4
0 Mbp
10 Mbp
20 Mbp
30 Mbp
40 Mbp
50 Mbp
60 Mbp
Chr08
0.0
0.2
0.4
0 Mbp
10 Mbp
20 Mbp
30 Mbp
40 Mbp
50 Mbp
60 Mbp
70 Mbp
Chr09
0.0
0.1
0.2
0.3
0 Mbp
5 Mbp
10 Mbp
15 Mbp
20 Mbp
25 Mbp
30 Mbp
35 Mbp
40 Mbp

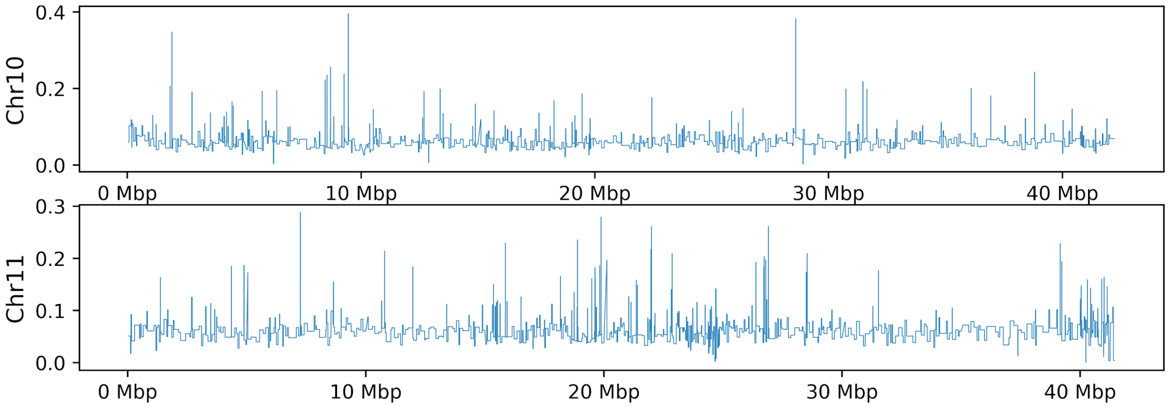


**Supplementary Figure S7. *E. sideroxylon* recombination**

| | | *E. melliodora* | *E.sideroxylon* |
|---|---|---|---|
| **Syntenic** | **Gene** | 0 | 0 |
| | **Transposon** | 0 | 0 |
| | **Inverted** | 0.9982 | 0.9942 |
| | **Translocated** | 1 | 0.5106 |
| | **Duplicated** | 0.0023 | 0.0151 |
| | **Unaligned** | 1 | 0.9969 |
| **Gene** | **Syntenic** | 0 | 0 |
| | **Transposon** | 0 | 0 |
| | **Inverted** | 0.9504 | 0.5622 |
| | **Translocated** | 0 | 0 |
| | **Duplicated** | 0 | 0 |
| | **Unaligned** | 0 | 0 |
| **Transposon** | **Syntenic** | 0 | 0 |
| | **Gene** | 0 | 0 |
| | **Inverted** | 0.9997 | 0.9981 |
| | **Translocated** | 0 | 0.002 |
| | **Duplicated** | 0 | 0 |
| | **Unaligned** | 0 | 0 |
| **Inverted** | **Syntenic** | 0.9982 | 0.9942 |
| | **Gene** | 0.9504 | 0.5622 |
| | **Transposon** | 0.9997 | 0.9981 |
| | **Translocated** | 0.9983 | 1 |
| | **Duplicated** | 1 | 1 |
| | **Unaligned** | 0.9977 | 0.9969 |
| **Translocated** | **Syntenic** | 1 | 0.5106 |
| | **Gene** | 0 | 0 |
| | **Transposon** | 0 | 0.002 |
| | **Inverted** | 0.9983 | 1 |
| | **Duplicated** | 0.699 | 1 |
| | **Unaligned** | 1 | 0.801 |
| **Duplicated** | **Syntenic** | 0.0023 | 0.0151 |
| | **Gene** | 0 | 0 |
| | **Transposon** | 0 | 0 |
| | **Inverted** | 1 | 1 |
| | **Translocated** | 0.699 | 1 |
| | **Unaligned** | 0.0138 | 0.3032 |
| **Unaligned** | **Syntenic** | 1 | 0.9969 |
| | **Gene** | 0 | 0 |
| | **Transposon** | 0 | 0 |
| | **Inverted** | 0.9977 | 0.9969 |
| | **Translocated** | 1 | 0.801 |
| | **Duplicated** | 0.0138 | 0.3032 |

**Supplementary Table S3. Pairwise rho Tukey's test p-values.** Green indicates a significant difference (P<=0.05).

| | | *E. melliodora* | *E.sideroxylon* |
|---|---|---|---|
| **Syntenic** | **Gene** | 0 | 0 |
| | **Transposon** | 0 | 0 |
| | **Inverted** | 0.9996 | 0.999 |
| | **Translocated** | 0 | 0 |
| | **Duplicated** | 0 | 0 |
| | **Unaligned** | 0 | 0 |
| **Gene** | **Syntenic** | 0 | 0 |
| | **Transposon** | 0 | 0 |
| | **Inverted** | 1 | 0.9605 |
| | **Translocated** | 0 | 0 |
| | **Duplicated** | 0 | 0 |
| | **Unaligned** | 0 | 0 |
| **Transposon** | **Syntenic** | 0 | 0 |
| | **Gene** | 0 | 0 |
| | **Inverted** | 0.8074 | 0.8693 |
| | **Translocated** | 0.1234 | 0.9997 |
| | **Duplicated** | 0.9983 | 0 |
| | **Unaligned** | 1 | 0 |
| **Inverted** | **Syntenic** | 0.9996 | 0.999 |
| | **Gene** | 1 | 0.9605 |
| | **Transposon** | 0.8074 | 0.8693 |
| | **Translocated** | 0.5745 | 0.8445 |
| | **Duplicated** | 0.7877 | 0.9975 |
| | **Unaligned** | 0.8103 | 1 |
| **Translocated** | **Syntenic** | 0 | 0 |
| | **Gene** | 0 | 0 |
| | **Transposon** | 0.1234 | 0.9997 |
| | **Inverted** | 0.5745 | 0.8445 |
| | **Duplicated** | 0.2856 | 0.004 |
| | **Unaligned** | 0.1464 | 0 |
| **Duplicated** | **Syntenic** | 0 | 0 |
| | **Gene** | 0 | 0 |
| | **Transposon** | 0.9983 | 0 |
| | **Inverted** | 0.7877 | 0.9975 |
| | **Translocated** | 0.2856 | 0.004 |
| | **Unaligned** | 0.9987 | 0.2449 |
| **Unaligned** | **Syntenic** | 0 | 0 |
| | **Gene** | 0 | 0 |
| | **Transposon** | 1 | 0 |
| | **Inverted** | 0.8103 | 1 |
| | **Translocated** | 0.1464 | 0 |
| | **Duplicated** | 0.9987 | 0.2449 |

**Supplementary Table S4. Pairwise Fst Tukey's test p-values.** Green indicates a significant difference (P<=0.05).

| | | *E. melliodora* | *E.sideroxylon* |
|---|---|---|---|
| **Syntenic** | **Gene** | 0 | 0 |
| | **Transposon** | 0 | 0 |
| | **Inverted** | 0.9349 | 0.4655 |
| | **Translocated** | 0 | 0.0029 |
| | **Duplicated** | 0 | 0 |
| | **Unaligned** | 0 | 0 |
| **Gene** | **Syntenic** | 0 | 0 |
| | **Transposon** | 0 | 0 |
| | **Inverted** | 0.4392 | 0.3614 |
| | **Translocated** | 0 | 0 |
| | **Duplicated** | 0.9991 | 0 |
| | **Unaligned** | 0 | 0 |
| **Transposon** | **Syntenic** | 0 | 0 |
| | **Gene** | 0 | 0 |
| | **Inverted** | 1 | 0.9755 |
| | **Translocated** | 0.0107 | 0.493 |
| | **Duplicated** | 0 | 0 |
| | **Unaligned** | 0 | 0 |
| **Inverted** | **Syntenic** | 0.9349 | 0.4655 |
| | **Gene** | 0.4392 | 0.3614 |
| | **Transposon** | 1 | 0.9755 |
| | **Translocated** | 0.9895 | 0.8918 |
| | **Duplicated** | 0.4565 | 1 |
| | **Unaligned** | 0.4602 | 0.0158 |
| **Translocated** | **Syntenic** | 0 | 0.0029 |
| | **Gene** | 0 | 0 |
| | **Transposon** | 0.0107 | 0.493 |
| | **Inverted** | 0.9895 | 0.8918 |
| | **Duplicated** | 0 | 0 |
| | **Unaligned** | 0 | 0 |
| **Duplicated** | **Syntenic** | 0 | 0 |
| | **Gene** | 0.9991 | 0 |
| | **Transposon** | 0 | 0 |
| | **Inverted** | 0.4565 | 1 |
| | **Translocated** | 0 | 0 |
| | **Unaligned** | 0 | 0 |
| **Unaligned** | **Syntenic** | 0 | 0 |
| | **Gene** | 0 | 0 |
| | **Transposon** | 0 | 0 |
| | **Inverted** | 0.4602 | 0.0158 |
| | **Translocated** | 0 | 0 |
| | **Duplicated** | 0 | 0 |

**Supplementary Table S5. Pairwise SNP per Kilobase Tukey's test p-values.** Green indicates a significant difference (P<=0.05).
